# A systems-level insight into PHB-driven metabolic adaptation orchestrated by the PHB-binding transcriptional regulator AniA (PhaR)

**DOI:** 10.1101/2025.02.17.638283

**Authors:** Antonio Lagares, Elizaveta Krol, Tina Müller, Timo Glatter, Anke Becker

## Abstract

Polyhydroxybutyrate (PHB) is a carbon and energy storage polymer, whose accumulation under nutrient imbalances with excess carbon is a widespread phenomenon in bacteria. PhaR is a conserved transcriptional regulator found to be associated to PHB granules in several species. Although its role in modulating PHB storage and metabolism has been extensively studied across the bacterial phylogeny, a comprehensive analysis of the PhaR regulon within the context of its dual role as a metabolic sensor and regulator remains missing. To bridge this gap, we integrated co-expression network analysis with proteome profiling across multiple mutant backgrounds (lack of PhaR (AniA) and/or PHB synthesis) in the free-living state of the PHB-accumulating alphaproteobacterial root nodule symbiont *Sinorhizobium meliloti*. This analysis was enriched by identifying direct regulatory targets of PhaR through a regulon-centric computational multi-step search for DNA binding site motifs combined with PhaR-DNA binding and promoter-reporter assays. We confirmed that the model of accumulated PHB sequestering PhaR and thereby relieving phasin and PHB depolymerase gene repression to control cellular PHB levels also applies to *S. meliloti*, and showed that PhaR-mediated regulation also occurs the symbiotic state. A comprehensive picture of the impact of PHB-mediated PhaR titration on cellular functions revealed exopolysaccharide production as well as central carbon metabolism (*pdh* and *bkd*), gluconeogenesis (*ppdK* and *pyc),* entry into the TCA cycle (*gltA*), and the initial steps of the Entner-Doudoroff pathway (*zwf, pgl and edd*) as major regulatory targets, and beyond carbon metabolism several target genes of yet unknown function. Our findings highlight a pivotal role for PhaR in orchestrating carbon metabolism.

## Introduction

Polyhydroxybutyrate (PHB) is a storage polymer accumulated by many bacterial species in the form of intracellular granules, called carbonosomes (Jendrossek, 2009; Jendrossek & Pfeiffer, 2014). Under growth-limiting conditions and if a utilizable C source is available, PHB is synthesized through the polymerization of beta-hydroxybutyryl residues by a PHB synthase. The process begins with the condensation of two acetyl-CoA molecules to form acetoacetyl-CoA, followed by reduction to beta-hydroxybutyryl-CoA by means of NAD(P)H. These steps are catalyzed by beta-ketothiolase and acetoacetyl-CoA reductase, respectively (Trainer & Charles, 2006). PHB is degraded by PHB depolymerases, and the products are modified yielding acetyl-CoA and reduced redox coenzymes. The PHB granule surface is shielded from the bacterial cytoplasm by phasins; other PHB granule-associated proteins include PHB synthases and depolymerases as well as a transcription regulator called PhaR in many species (Grage et al., 2003).

In the model organism for PHB biosynthesis, the betaproteobacterium *Cupriavidus necator* (formerly *Ralstonia eutropha*), PhaR has been shown to repress the transcription of genes encoding phasins (York et al., 2002). PhaR acts as a transcription repressor when no PHB is produced, and is sequestered to the surface of PHB granules away from the DNA when PHB accumulates, thus ensuring tight regulation of phasin abundance relative to produced PHB (Pötter et al., 2005a). Similar functions of PhaR homologs were also identified in alphaproteobacteria; in *Rhodobacter sphaeroides* and *Paracoccus denitrificans,* PhaR also represses phasin gene expression (Chou et al., 2009; Maehara et al., 2002). The ability of *R. spaeroides* PhaR to bind DNA was confirmed, and the binding sequence was determined as 5’-CTGCNNNGCAG-3’ (Chou & Yang, 2010). *In vitro* binding of PhaR to PHB was documented in *Paracoccus denitrificans* (Maehara et al., 2002). In the alphaproteobacterial rhizobia *Rhizobium etli* and *Bradyrhizobium diazoefficiens*, identified PhaR homologs were shown to have a broader role in gene expression control (Encarnación et al., 2002a; Nishihata et al., 2018; Quelas et al., 2016, 2024).

Despite these advances in understanding the function of PhaR, no study has yet provided a high-resolution global analysis of the PhaR regulon in the context of its dual role as a metabolic sensor and regulator. To fill this gap, we devised an approach combining co-expression network analysis with proteomics across multiple mutant backgrounds using the model symbiotic soil-dwelling α-proteobacterium *Sinorhizobium meliloti*. In *S. meliloti,* a chromosomally encoded *phaR* homolog was first identified and named *aniA* (for *a*naerobically *i*nduced *g*ene *A*; for consistency renamed *phaR* in our study) by Povolo and Casella (Povolo & Casella, 2000a). The symbiosis between *S. meliloti* and legumes of the genera *Melilotus*, *Medicago*, and *Trigonella* results in the formation of indeterminate root nodules, in which the colonizing bacteria exist either in a vegetative state or as differentiated nitrogen-fixing bacteroids. Establishment of symbiosis is a complex multistep process that involves signaling and adaptation. The extensive genetic, phenotypic, and molecular characterization of *S. meliloti*, combined with an advanced global understanding of its metabolism in its free-living and symbiotic lifestyles, makes it an ideal model for exploring the dual role of PhaR (AniA) at a molecular systems-level (diCenzo et al., 2019).

*S. meliloti* stores excess carbon as intracellular PHB granules as its main carbon storage compound (Zevenhuizen, 1981), and its PHB metabolon has been extensively studied (Charles et al., 1997; Tombolini et al., 1995; Trainer et al., 2010; Willis & Walker, 1998). The phasins PhaP1 (SMc00777) and PhaP2 (SMc02111) were identified as granule-associated proteins necessary for efficient PHB accumulation (C. Wang, Sheng, et al., 2007). An active PHB metabolism in free-living conditions has been shown to play a crucial role in regulating the expression of nitrogen metabolism genes, likely through the involvement of an Fnr-type transcription factor (Alessio et al., 2017). Studies by Ratcliff and Denison (Ratcliff et al., 2008; Ratcliff & Denison, 2010, 2011) explored PHB’s role in bet-hedging and persistence, demonstrating asymmetrical distribution of PHB granules between mother and daughter cells under nutrient limitation in *S. meliloti*. In symbiosis, PHB is accumulated at early stages during the infection, but PHB granules are not observed in N_2_-fixing bacteroids (Trainer & Charles, 2006).

*S. meliloti* PhaR shares between 39% and 83% amino acid similarity with studied PhaR homologs from alpha- and beta-proteobacteria. Mutation of *phaR* (*aniA*) in *S. meliloti* Rm41 reduces PHB accumulation while increasing glycogen and EPS production. Furthermore, this mutation alleviates growth defects in a *phbC* mutant when pyruvate is used as the sole carbon source (Povolo & Casella, 2000b, 2009a). Similar findings in *R. etli phaR* (*aniA*) mutants confirm the conserved function of this gene in symbiotic rhizobia (Encarnación et al., 2002b). Moreover, *phaR* depletion in *S. meliloti* leads to overproduction of the small RNA MmgR (Ceizel Borella et al., 2018), which finetunes PHB accumulation, likely via post-transcriptional repression of phasin gene expression (Lagares, Borella, et al., 2017). Consequently, functional PhaR may be crucial for proper phasin production during the onset of PHB synthesis. While the ability to produce PHB was not crucial for symbiotic performance of *S. meliloti*, PhaR provided a competitive advantage at early stages of the symbiotic interaction (Aneja et al., 2005). Along with PHB, *S. meliloti* accumulates glycogen and produces the exopolysaccharides succinoglycan (EPSI) and galactoglucan (EPSII), which have a critical role during symbiosis (Jones et al., 2007).

Our study provides broad and deep insights into the PhaR regulon in *S. meliloti* through a systems-level analysis. We advance the knowledge of the PhaR DNA binding site by uncovering previously undetected extended AT-rich arms, and describe conserved PhaR regulatory targets in the PHB metabolon, novel targets that are likely conserved but not yet reported in PhaR-bearing bacteria, and variations in targets among phylogenetically related bacteria. We report proteomic, biochemical, and phenotypic evidence supporting a role for PhaR in regulating key steps of central carbon and EPS metabolism consistent with its previously proposed role in coordinating carbon fate under nutrient imbalance, and implying a broader regulatory role beyond intermediate metabolism. Our co-expression network analysis reveals that PhaR functions as both a metabolic sensor and a global gene regulator by combining its evolutionarily conserved ability to bind both DNA and PHB.

## Material and Methods

### Strains, plasmids and growth conditions

The strains and plasmids used in this study are listed in the Table S1. *E. coli* used for cloning and conjugation were grown at 37°C on LB medium (10 g/L Tryptone, 5 g/L Yeast extract, 5 g/L NaCl). When required, kanamycin (50 mg/L) or tetracycline (10 mg/L) was added. *S. meliloti* was cultured at 30°C on TY medium (5 g/L Tryptone, 3 g/L Yeast extract, 0.4 g/L CaCl_2_.2H_2_O), or MOPS-buffered phosphate-limiting minimal medium, regarded further as low P medium (10 g/L MOPS, 10 g/L Mannitol, 3,55 g/L Na glutamate, 1 mM MgSO_4_, 0.25 mM CaCl_2_, 10mg/L FeCl_3_.6H_2_O, 1 mg/L biotin, 3 mg/L H_3_BO_3_, 2.23 mg/L MnSO_4_.4H_2_O, 0.287 mg/L ZnSO_4_.7H_2_O, 0.125 mg/L CuSO_4_.5H_2_O, 0.065 mg/L CoCl_2_.6 H_2_O, 0.12 mg/L NaMoO_4_.2H_2_O, 0.1 mM KH_2_PO_4_). When required, streptomycin was added at 600 mg/L, kanamycin at 200 mg/L and tetracycline at 10 mg/L. In liquid cultures, antibiotic concentrations were halved.

### Cloning and genetic manipulations

The constructs used in this work were generated using standard genetic techniques. Plasmids were transferred to *S. meliloti* by conjugation with *E. coli* S17-1. The primers used are listed in the Table S1.

Deletions of *phaR* and *phbC* were generated using constructs pPhaRdel and pPhbCdel, which contained DNA fragments upstream and downstream of the target gene, along with the negative sucrose selection marker *sacB*. Double recombinants with gene deletions were selected on LB agar containing 10% sucrose, as previously described (Lagares et al., 2017), and verified by PCR.

To generate C-terminal translation fusions to EGFP at the native genomic locus, the 3’-terminal portion of the gene (approximately 300 bp, excluding the stop codon) was cloned into the suicide vector pK18mob2-EGFP to generate an in-frame fusion. The resulting constructs were integrated into the *S. meliloti* genome.

Promoter-*EGFP* fusions were generated by inserting the promoter region, including the translation start codon and up to the first 10 codons of the downstream gene, into the replicative low-copy plasmid pPHU231-EGFP or the replicative medium-copy plasmids pSRKKm-EGFP or pBBRKm-EGFP. This created an in-frame fusion of the first few codons of the gene of interest to *EGFP*.

A construct for the purification of His_6_-tagged PhaR was generated by inserting the protein coding sequence into the expression vector pWH844.

### EGFP measurements

For promoter-*EGFP* assays, overnight precultures in TY medium were diluted 1:500 in fresh culture medium and grown in a 96-well polystyrene flat-bottom plate (Greiner) with 100 µl of medium per well. The plates were shaken at 1200 rpm at 30°C. EGFP fluorescence and optical density were measured using a TECAN spectrophotometer (excitation 488 nm, emission 522 nm, gain 82). Fluorescence values were calculated as relative fluorescence units (RFU), which represent the fluorescence intensity divided by the optical density. Unless otherwise indicated, background fluorescence from corresponding strains carrying empty vectors pPHU231-EGFP or pSRKKm-EGFP was subtracted. Three to four independent transconjugants of each strain containing the promoter-EGFP fusion were used as biological replicates.

### Protein purification

*E. coli* BL21 strain carrying pWH844-*phaR* was grown in LB with ampicillin (100 µg/ml) to an optical density at 600 nm of 0.4 at 37^°^C, then induced with 500 µM IPTG for 7 hours at 30^°^C or for 16 h at room temperature. Cells from 100 ml of culture were harvested by centrifugation at 4,000 x*g*, resuspended in 6 ml of binding buffer (1.76 g/L Na_2_HPO_4_.2H_2_O, 1.4 g/L NaH_2_PO_4_.H_2_O, 29.2 g/L NaCl, 20 mM imidazole, pH 7.4) and lysed using a French press. The lysates were cleared by centrifugation at 20,000 x*g* and 4°C for 40 min, and 2400 µl of the supernatant was applied to a His SpinTrap column (GE Healthcare). The column was washed 3 times with binding buffer, and the protein was eluted with elution buffer (binding buffer supplemented with 500 mM imidazole). Protein purity was assessed by SDS-PAGE and Coomassie blue staining, and the protein concentration was determined using the Bradford reagent.

### *In vitro* PHB binding

The binding reactions were performed using 10 µg of protein and 0, 0.1, 0.25 or 1 µg of crystalline PHB in 5 mM Tris-HCl (pH 8.5), in a total volume of 100 µl, in triplicate. Reactions were incubated at room temperature (RT) for 1 hour with shaking at 1000 rpm, then centrifuged at 20,000 x *g* for 5 minutes at RT. Ninety microliters of the supernatant were mixed with 410 µl of water and subjected to a Bradford assay to estimate the remaining soluble protein.

### Electrophoretic mobility shift assay (EMSA)

An EMSA reaction mixture consisted of 10 mM Tris-HCl (pH 8.5), 50 mM KCl, 0.025 *A*260 units of sonicated salmon sperm DNA (GE Healthcare), 1.0 mg/ml of bovine serum albumin (Sigma), and 10 ng of Cy3-labelled DNA, in a final volume of 10 µl. The protein was added as indicated, in 1 µl of an appropriate dilution in the elution buffer. The reaction mixture was incubated at room temperature for 30 min. Then, 1.5 µl of 90% glycerol was added, and the reaction mixtures were loaded onto a 2% agarose gel. Following electrophoresis at 3 V/cm in TAE buffer (40mM Tris, 20 mM acetic acid, 1 mM EDTA) at room temperature for 2 h, gels were scanned using a Typhoon 8600 variable-mode imager (Amersham Bioscience).

### Plant assays

*Medicago sativa* cv. Eugenia seeds were surface-sterilized with sulfuric acid for 10 min, washed six times with sterile water and transferred to plant agar (Broughton & Dilworth, 1971). The seeds were germinated in the dark overnight at 4 °C, followed by 24 h at 30 °C. Bacterial overnight cultures in TY medium were diluted 1:10 with sterile water, and 100 μl of the suspension was plated onto the lower half of plant agar plates. Four seedlings were placed on each plate, and plants were grown at 22 °C and 90 % humidity in a 18 h light / 6 h night photoperiod for three weeks.

To determine shoot dry weight, the shoots of four plants from a single agar plate were dried at 60°C for two days and then weighted. Nodule numbers were assessed by visual examination. The total number of nodules per plate (four plants) was counted and used to calculate the average number of nodules per plant.

### Microscopy

Microscopy of bacteria on 1% agarose pads was performed using Nikon Eclipse Ti-E microscope with DIC objective 100x, CFI Apochromat TIRF Oil objective (numerical aperture of 1.49) and phase contrast objective Plan Apo l 100×1.45 oil with AHF HC filter sets F36-504 TxRed (ex bp 562/40 nm, bs 593 nm, em bp 624/40 nm) and F36-525 EGFP (exc bp 472/30 nm, beam splitter 495 nm, em bp 520/35 nm). Images were acquired with an Andor iXon3 885 EMCCD camera.

For time-lapse microscopy, bacteria were imaged on low phosphate medium agarose pads at 30^°^C, with images acquired every 15 min.

Microscopy of nodules was performed on fresh 100 µm longitudal nodule sections using a 20x objective lens. Nodule slices were prepared using a Leica VT 1000 S vibratome.

### Total protein quantification

Total protein content was determined with the Bicinchoninic Acid (BCA) assay (Pierce BCA Protein Assay Kit, Thermo Scientific). Briefly, biomass equivalent to 1 OD unit was collected from each culture by centrifugation and resuspended in 100 µl of Laemmli buffer lacking bromophenol blue (composition). The cells were lysed by heating at 100°C for 10 minutes, followed by centrifugation at 12,000xg for 1 min. Protein quantification was performed on 10 µl of the clarified supernatant according to the manufacturer’s instructions. A standard curve was generated using Bovine Serum Albumin (BSA) solutions prepared in the same buffer, which was used to interpolate protein concentrations in the samples.

### PHB quantification

Nile red–based staining was employed to visualize PHB granules using a fluorescence microscope and to quantify polyhydroxybutyrate (PHB) in stationary-phase cultures of *S. meliloti* strains under different growth conditions, following a previously published protocol (Lagares Jr. & Valverde, 2017). Briefly, biomass equivalent to 0.5 OD units was collected by centrifugation (1 min, 12,000 x*g*) and incubated for 30 min with 35% EtOH in PBS (pH 7.2). The cells were centrifuged (1 min, 12,000x*g*), the supernatant was discarded, and the pellet was resuspended in 1 ml PBS (pH 7.2). From a 0.1 mg/mL Nile Red stock solution in DMSO, 10 μL was added to the bacterial suspension. Following 30 min of incubation, 100 μL aliquots were transferred to a 96-well plate, and optical density (OD, 600 nm) and fluorescence (excitation at 488 nm, emission at 585 nm) were measured using a Tecan Infinity plate reader. Fluorescence values were normalized to OD, and subsequently to total protein content, yielding arbitrary fluorescence units (AFU) per µg protein.

### Quantitative proteomic analyses

A bacterial biomass from 1 ml of cultures that were grown for 48 h in TY and MOPS-buffered low P media was collected by centrifugation at 10,000 × *g* for 3 min at RT. The pellets were resuspended in 150 µl of milliQ water and transferred to a screw-cap 2-ml tube filled with ca. 70 mg of 0.1 mm glass beads. Bacterial cells were lysed by four 60-sec-cycles of homogenization at 6.5 m/s in a FastPrep-24 device (MP Biomedicals). Then, 150 µl of a 2× lysis buffer (40 mM HEPES pH 7.5, 4% Sodium Lauryl Sarcosinate) was added to the suspensions and the samples were further incubated at 90°C for 15 min. The lysates were clarified by centrifugation at 20,000 × *g* for 5 min at RT, and the supernatants were stored at -20°C. The total protein content in the lysates was quantified with the Bicinchoninic Acid method (BCA Protein Assay Kit), and a sample volume containing 50 µg of protein was submitted for proteomic analysis.

Proteins were reduced with 5 mM Tris(2-carboxyethyl) phosphine (Thermo Fischer Scientific) at 90°C for 15 min and alkylated using 10 mM iodoacetamid (Sigma Aldrich) at 20°C for 30 min in the dark. Proteins were precipitated with a 6-fold excess of ice-cold acetone, followed by two methanol washing steps. Dried proteins were reconstituted in 0.2 % SLS and the amount of proteins was determined by bicinchoninic acid protein assay (Thermo Scientific). For tryptic digestion 50 μg protein was incubated in 0.5% SLS and 1 μg of trypsin (Serva) at 30°C overnight. After digestion, SLS was precipitated by adding a final concentration of 1.5% trifluoroacetic acid (TFA, Thermo Fischer Scientific). Peptides were desalted by using C18 solid phase extraction cartridges (Macherey-Nagel). Cartridges were prepared by adding acetonitrile (ACN), followed by equilibration with 0.1% TFA. Peptides were loaded on equilibrated cartridges, washed with 5% ACN and 0.1% TFA containing buffer and finally eluted with 50% ACN and 0.1% TFA. Peptides were dried and reconstituted in 0.1% trifluoroacetic acid and then analyzed using liquid-chromatography-mass spectrometry carried out on a Exploris 480 instrument connected to an Ultimate 3000 RSLC nano and a nanospray flex ion source (all Thermo Scientific). Peptide separation was performed on a reverse phase HPLC column (75 μm x 42 cm) packed in-house with C18 resin (2.4 μm; Dr. Maisch). The following separating gradient was used: 94% solvent A (0.15% formic acid) and 6% solvent B (99.85% acetonitrile, 0.15% formic acid) to 25% solvent B over 95 minutes at a flow rate of 300 nl/min, and an additional increase of solvent B to 35% for 25 min. MS raw data was acquired in data independent acquisition mode with a method adopted from Bekker-Jensen et al. (Bekker-Jensen et al., 2020). In short, Spray voltage were set to 2.3 kV, funnel RF level at 40, and heated capillary temperature at 275 °C. For DIA experiments full MS resolutions were set to 120.000 at m/z 200 and full MS, AGC (Automatic Gain Control) target was 300% with an IT of 50 ms. Mass range was set to 350–1400. AGC target value for fragment spectra was set at 3000%. 45 windows of 14 Da were used with an overlap of 1 Da. Resolution was set to 15,000 and IT to 22 ms. Stepped HCD collision energy of 25, 27.5, 30 % was used. MS1 data was acquired in profile, MS2 DIA data in centroid mode.

Analysis of DIA data was performed using DIA-NN version 1.8 using a UniProt protein database from *Sinorhizobium meliloti* 1021 (Consortium, 2023; Demichev et al., 2020). Full tryptic digest was allowed with two missed cleavage sites, and oxidized methionines and carbamidomethylated cysteine. Match between runs and remove likely interferences were enabled. The neural network classifier was set to the single-pass mode, and protein inference was based on genes. Quantification strategy was set to any LC (high accuracy). Cross-run normalisation was set to RT-dependent. Library generation was set to smart profiling. DIA-NN outputs were further evaluated using the SafeQuant and script modified to process DIA-NN outputs (Ahrné et al., 2013; Glatter et al., 2012). The SafeQuant script was executed on the “report.tsv” file from DIA-NN analysis to sum precursor intensities to represent protein intensities. The peptide-to-protein assignment was done in SafeQuant with redundant peptide assignment following the Occam’s razor approach. Median protein intensity normalization was performed followed by imputation of missing values using a normal distribution function.

The statistical analysis was performed in Perseus (Tyanova & Cox, 2018) and R (Team, 2014). Proteins with at least two unique peptides identified were considered for further analysis. To ensure robust detection, a protein was retained if at least two unique peptides were detected in at least two out of three biological replicates for at least two out of the four mutant strains. Missing values for proteins meeting these criteria were imputed using a Gaussian distribution with a standard deviation of 0.3 and a downshift of 1.8 standard deviations, with these parameters set separately for each sample’s proteome. ANOVA analyses of proteomes from cells grown in low P or TY medium were performed in Perseus. The results were filtered using a permutation-based false discovery rate (FDR) correction, with a significance threshold for q-values set at 0.05. Z-score data normalization was carried out in Perseus accessing the matrix by rows with grouping set by growth medium. Principal component analysis (PCA) was performed using the Factoextra package for R (Kassambara & Mundt, 2020). Z-scored data was used to run a co-expression network analysis with the hCoCena package for R (Oestreich et al., 2022), following the protocol recently published by the developers (Holsten et al., 2024).

## Results

### PhaR binds PHB *in vitro* and regulates global gene expression dependent on PHB production

The ability of PhaR from *S. meliloti* to bind PHB was evaluated *in vitro*. Purified His_6_-PhaR, but not BSA which was conceived as a negative control, was effectively removed from the solution upon incubation with a suspension of non-soluble PHB and subsequent centrifugation (Fig. 1A). We determined that approx. 10 µg of His_6_-PhaR (ca. 440 pmoles) bind to 1 µg of crystalline PHB in a total volume of 100 µl (Fig. 1A).

**Figure 1.**
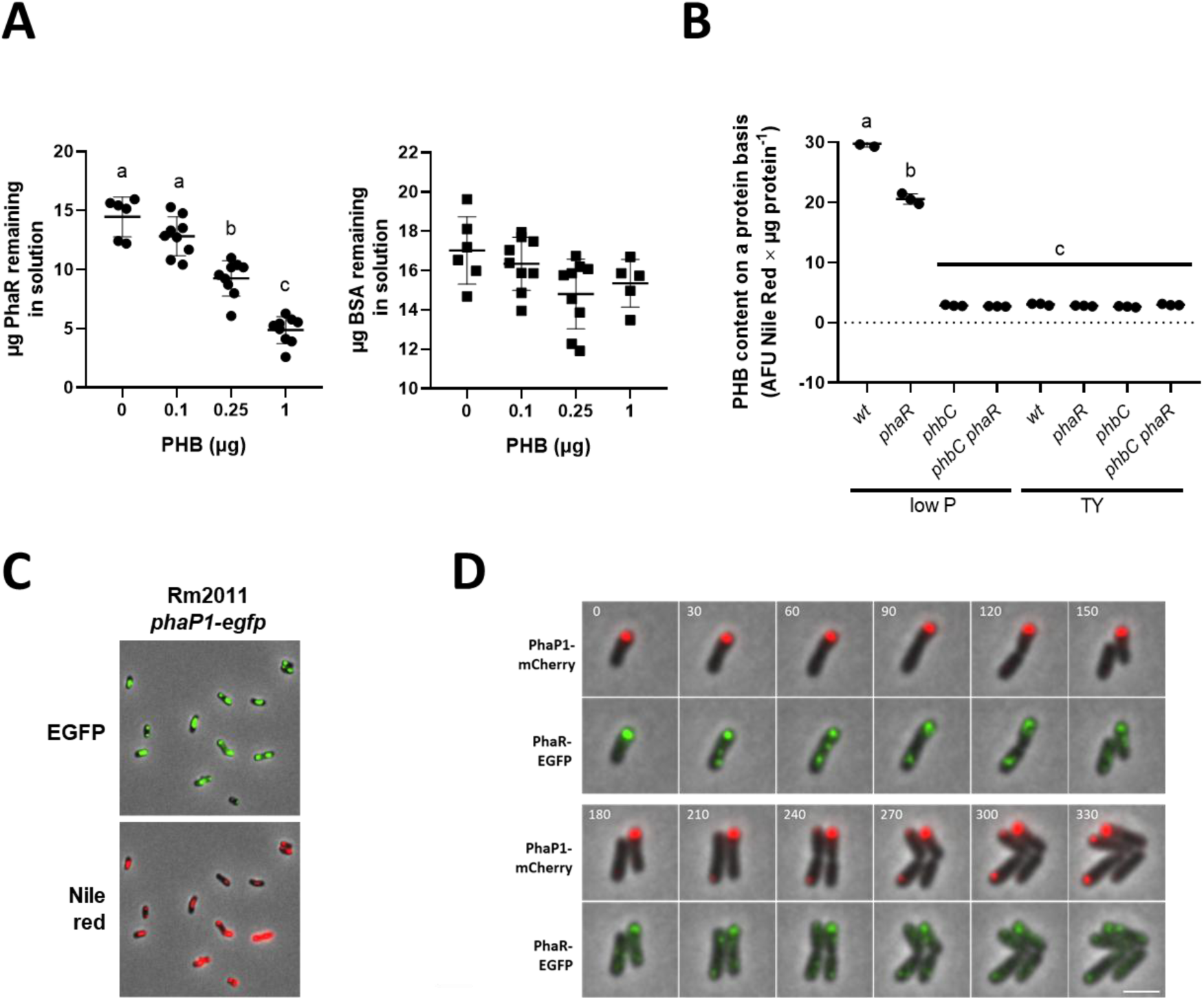
PhaR associates with PHB granules and modulates PHB accumulation under nutrient-limiting conditions in low-phosphate medium. **(A)** *In vitro* binding of crystalline PHB by His_6_-PhaR. The total protein remaining in solution after incubation of PhaR (filled circles) or BSA (filled squares) with increasing amounts of crystalline PHB is shown. Error bars represent the standard deviation from at least five replicates. Statistical analysis for each experiment was conducted using ANOVA, followed by Tukey’s multiple-comparison test. Bars labeled with different letters indicate statistically significant differences (P< 0.001). **(B)** PHB levels to total protein are shown with filled circles for wild type (wt), *phaR*, *phbC*, and *phbC phaR* strains in the stationary phase of growth under conditions permissive (low P medium) and non-permissive (TY medium) for PHB accumulation. **(C)** Microscopy images of cells stained with Nile red after 24 hours of growth in low P medium. Scale bar: 5 µm.

To obtain further evidence supporting the hypothesis that PhaR associates with PHB *in vivo*, we analyzed the colocalisation of phasins and PhaR with PHB granules. Phasin PhaP1, previously identified as a PHB granule-associated protein (C. Wang, Sheng, et al., 2007), was initially used to visualize PHB granules. For this purpose, the coding sequence of EGFP was integrated at the native genomic location of *phaP1* to replace its stop codon. As expected, PhaP1-EGFP foci colocalised with Nile Red-stained PHB granules (Fig. 1C), showing that this fusion protein can be used as marker for PHB granules. Colocalisation of PhaR-EGFP and PhaP1-mCherry observed by time-lapse microscopy in living cells then confirmed the recruitment of PhaR to PHB (Fig. 1D).

To characterize PhaR-mediated gene regulation and its dependence on PHB accumulation in *S. meliloti*, we generated Sm2011 mutants with deletions of *phaR* and/or PHB synthase gene *phbC*. PHB content was quantified in the wild type, as well as in the *phaR* and/or *phbC* mutant strains which were grown to the stationary phase in TY complex medium and MOPS-buffered minimal medium limited in phosphate, hereafter referred to as low P medium (Fig. 1B) (Krol & Becker, 2004). Consistent with previous reports (Povolo & Casella, 2000a), *phaR* cells grown in low P medium contained ∼30% less PHB than the wild type, while wild type and *phaR* cells grown in TY medium were indistinguishable from *phbC* cells that are unable to synthesize the carbon storage polymer. In agreement with these results, no PHB granules were visualized under the microscope upon Nile red staining of wild type or *phaR* cells grown in TY medium, or *phbC* mutant cells grown in either medium (Fig. S1). Thus, we further considered the growth in low P medium as permissive for PHB accumulation, in contrast to growth in TY medium, which supported no or little PHB production.

Sm2011 *phaR*, *phbC*, and *phbC phaR* strains formed pink, healthy nodules on *M. sativa* roots, indistinguishable from those induced by the wild type. Plant shoot dry weight and nodule numbers did not significantly differ between strains (Fig. 2A, B). We analyzed nodules from wild-type and *phaR* strains carrying chromosomally integrated phasin-EGFP fusions, using fluorescence as a readout of PhaR activity. Weak PhaP1-EGFP and PhaP2-EGFP signals appeared in the nitrogen fixation zone of wild-type nodules, while *phaR* mutants strongly overproduced PhaP1-EGFP, with PhaP2-EGFP unchanged (Fig. 2C). This mirrored the TY medium pattern (Fig. S1), which aligns with the fact that indeterminate nodule bacteria also produce little PHB, keeping PhaR free to regulate its target regulon.

**Figure 2.**
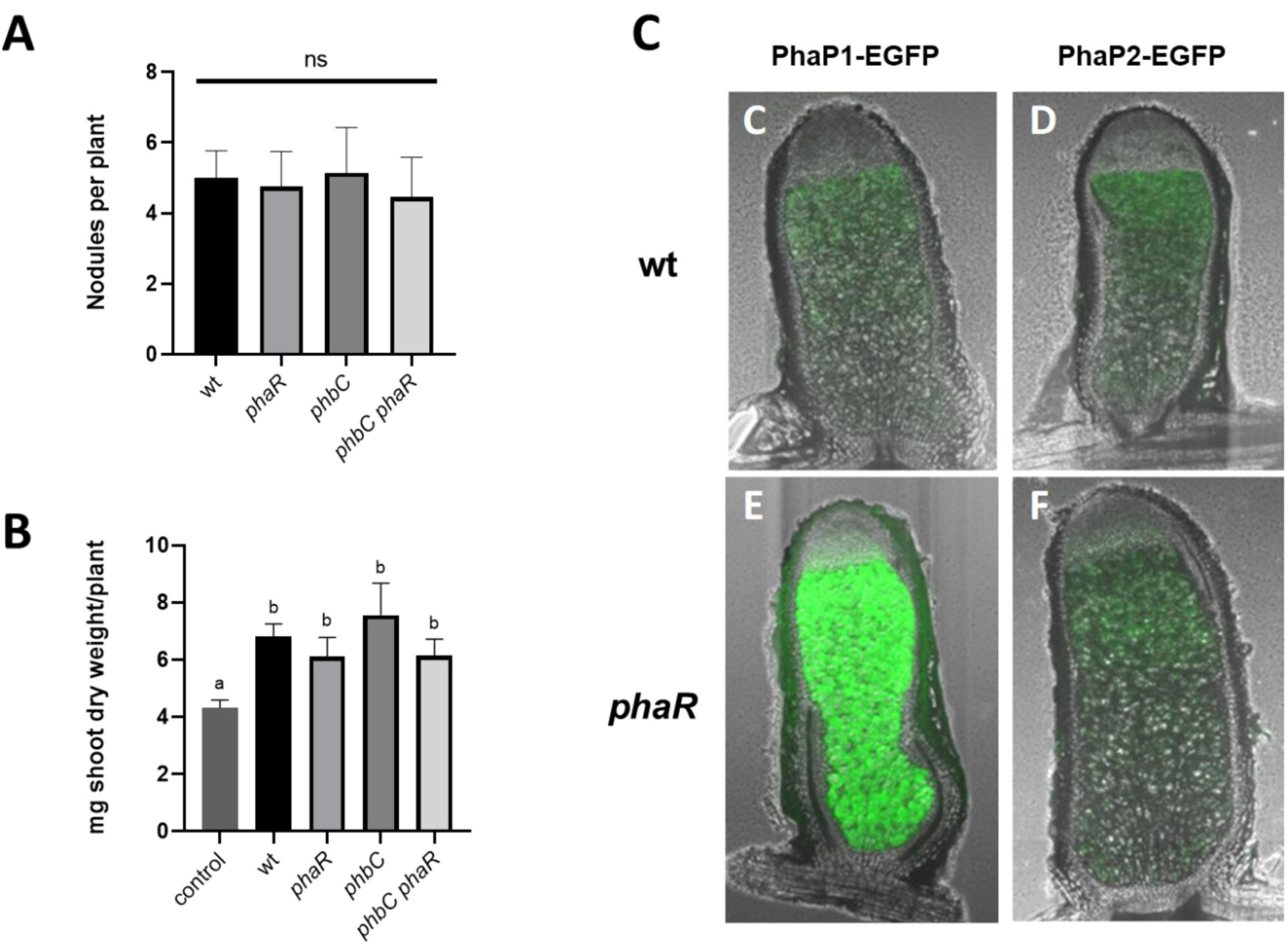
PhaR is active during symbiosis but is not essential for nodule formation or nitrogen fixation. **(A, B).** Symbiotic phenotypes of strains with varying PhaR and PhbC activity: **(A)** number of pink nodules formed per plant, (**B)** plant shoot dry weight. Data represent the averages from 20 plants inoculated with each of the indicated strains. Statistical analysis was performed using ANOVA followed by Tukey’s multiple-comparison test. Bars marked with different letters indicate significant differences (P < 0.01). **(C-F)** Microscopy images of longitudinal sections of nodules. **(C)** Sm2011 carrying P*_phaP1_*-*EGFP*, **(D)** Sm2011 carrying P*_phaP2_*-*EGFP*, **(E)** *phaR* mutant carrying P*_phaP1_*-*EGFP*, **(F)** *phaR* mutant carrying P*_phaP2_*-*EGFP*. Acquisition time was set at 1 s, except for E, for which it was set at 200 ms due to higher signal intensity.

Consistent with the model that PhaR homologs regulate transcription and are titrated by PHB granules, deletion of *phaR* significantly affected the proteome of *S. meliloti* only when PHB storage was minimal or absent (e.g., in *phbC* mutant cells grown in either medium, or in *phbC*^+^ cells grown in TY medium), as shown by a Principal Component Analysis (PCA) of the proteomic data (Fig. 3A). Furhermore, deletion of *phbC* under PHB-permissive conditions resulted in proteomic changes that contrasted with the effects of *phaR* deletion in a *phbC* mutant background. This difference is likely due to the release and increased availability of free PhaR molecules. With regard to the *phbC phaR* proteome, this is reflected in a nearly complete reversal of the shift of the *phbC* proteome relative to the wild type proteome along the x-axis of the PCA plot (Fig. 3A, left panel).

**Figure 3.**
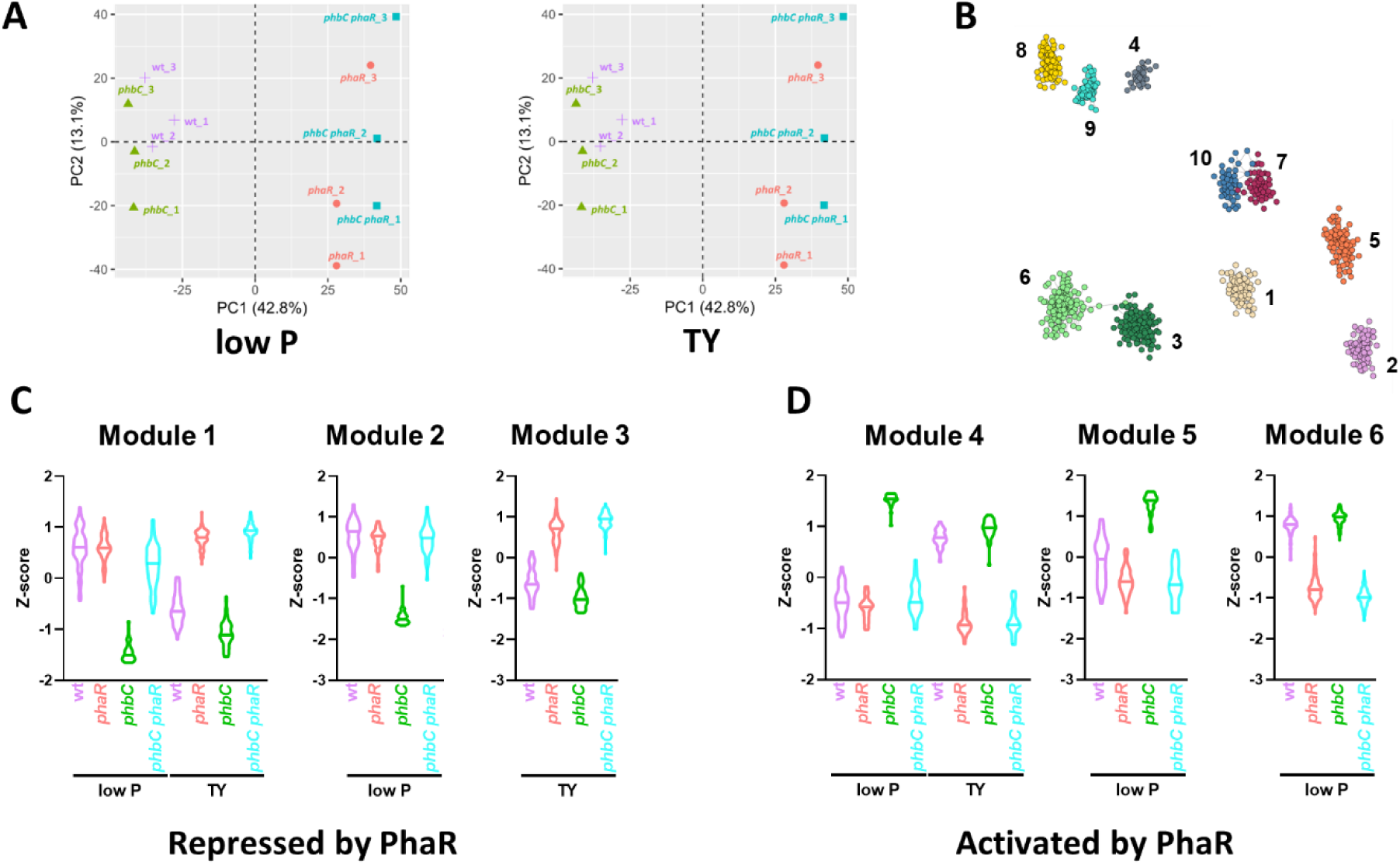
Co-expression networks reveal PhaR-driven proteome adjustments linked to PHB sensing. **(A)** Principal component analysis of proteomes from three biological replicates of wild type (purple symbols) and mutant (*phaR*, coral; *phbC*, green; *phbC phaR*, turquoise) strains grown in low P (left panel) and TY (right panel) media. **(B)** Network spatial visualization showing genes grouped based on their co-expression patterns across strains and growth conditions, as determined by the analysis using the hCoCena package. Genes are color-coded according to their respective co-expression modules, with each module assigned a signature number from 1 to 10. (**C)** Violin plot displaying the expression profiles of proteins repressed by PhaR, clustered in Modules 1 to 3, across mutant backgrounds and growth media. For each protein, the Z-scored abundances from three replicate proteomes were averaged. Each violin represents the distribution of these averaged protein abundances. **(D)** Violin plot displaying the expression profiles of proteins activated by PhaR, clustered in Modules 4 to 6, across mutant backgrounds and growth media. For each protein, the Z-scored abundances from three replicate proteomes were averaged. Each violin represents the distribution of these averaged protein abundances.

Building on our basic analyses, which confirmed that in *S. meliloti* PhaR interacts with PHB granules and that this interaction likely affects the regulatory activity of PhaR, we sought to identify the genes regulated by PhaR. To achieve this, we integrated the global protein expression data collected across various mutant backgrounds and growth conditions, and constructed a weighted co-expression network using the hoCoCena package to identify genes with expression patterns consistent with PhaR regulation (Oestreich et al., 2022). Genes with similar expression patterns across strains/conditions were clustered into modules. We identified ten modules that were further classified based on their enrichment in proteins of differential abundance in TY and low P media in the *phbC* strain compared to the *phbC phaR* strain (Fig. 3B). This pattern indicates proteins that are regulated by PhaR in a PHB-independent manner (Fig. 3C, D).

Modules 1 to 3 were enriched in proteins showing increased levels upon *phaR* deletion. The abundance profiles of these proteins across the different mutational backgrounds and growth conditions tested would be consistent with repression of their genes by PhaR (Fig. 3C). This supports the model in which PhaR actively regulates gene expression when this regulator is not sequestered by PHB. Conversely, modules 4 to 6 showed enrichment in proteins downregulated upon *phaR* deletion. In the tested settings, these proteins showed abundance profiles that would be consistent with activation of their genes by PhaR (Fig 3D). Proteins in modules 1 and 4 were produced in both media, whereas those in modules 2 and 5, and 3 and 6, were exclusively detected in low P and TY media, respectively. Modules 7 to 10 contained proteins whose abundance was differently affected by PhaR in the different growth conditions (not shown), likely reflecting indirect responses to PhaR or regulation influenced by external factors specific to each condition.

Modules 1 to 3 were significantly enriched in proteins involved in C storage metabolism, specifically related to PHB and EPS biosynthesis (Table 2). Levels of PHB granule-associated phasins PhaP1 and PhaP2, along with the PHB depolymerase SMa1961 were reduced in the *phaR* mutant background. Proteins encoded by the EPSI and EPSII biosynthetic clusters (*exo/exs* and *wga*/*wge*, respectively) are associated with these modules. Module 1 includes enzymes from central carbon metabolic pathways such as the Entner-Doudoroff (6-phosphogluconolactonase, Pgl; and phosphogluconate dehydratase, Edd), as well as enzymes from the TCA cycle and glyoxylate shunt (citrate synthase, GltA; isocitrate lyase, SMc01727), and gluconeogenesis (pyruvate phosphate dikinase, PpdK; pyruvate carboxylase, Pyc). Inorganic pyrophosphosphatase (Ppa) and urease components UreA and UreB are also grouped in Module 1, along with Tam (putative trans-aconitate methyltransferase, SMc00225) and Dhe (putative (S)-2-haloacid dehalogenase, SMc00103). ExoF3 (Module 2) and SMb21248 (Module 1) belong to a Clr-regulated 30-kb gene cluster involved in polysaccharide metabolism (Zou et al., 2017). From Module 1, the proteins SMc03998, SMc02051, SMb20609 and SMb20727, all with unknown functions, exhibited notable fold changes in the *phbC phaR* compared to the *phbC* strain (Table 2). Notably, in addition to the EPSII biosynthesis gene cluster-related proteins, NodP (encoding subunit 2 of the sulfate adenylyltransferase, an enzyme involved in sulphate activation) stands out in Module 2 with a strong upregulation in the absence of PhaR.

**Table 1.** Strains and plasmids used in this work.

| Strain | Properties | Reference |
| --- | --- | --- |
| ***S. meliloti*** |  |  |
| Sm2011 | Wild type, Str<sup>r</sup> | (Meade & Signer, 1977) |
| Rm101 | Sm2011, *mucR*::Spec<sup>r</sup> | (Becker et al., 1997) |
| *phaR* | Sm2011 Δ*phaR*, markerless deletion | This work |
| *phbC* | Sm2011 Δ*phbC*, markerless deletion | This work |
| *phbC phaR* | Sm2011 Δ*phbC* Δ*phaR*, markerless deletions | This work |
| *mucR phaR* | Rm101 Δ*phaR*, markerless deletion | This work |
| *mucR phbC* | Rm101 Δ*phbC*, markerless deletion | This work |
| *mucR phbC phaR* | Rm101 Δ*phbC* Δ*phaR*, markerless deletions | This work |
| Sm2011 PhaP1-EGFP | Sm2011 *phaP1*::pK18mob2-EGFP | This work |
| Sm2011 PhaP2-EGFP | Sm2011 *phaP2*::pK18mob2-EGFP | This work |
| Sm2011 PhaR-EGFP | Sm2011 *phaR*::pK18mob2-EGFP | This work |
| ***E. coli*** |  |  |
| DH5a | F<sup>−</sup> *endA1 supE44 thi-1 l- recA1 gyrA96 relA1 deoR* D(*lacZYA-argF*) U169 | (Grant et al., 1990) |
| S17-1 | *E. coli* 294 *thi* RP4-2-Tc::Mu-Km::Tn7 integrated into the chromosome | (Simon et al., 1983) |
| BL21 (DE3) | *fhuA2 [lon] ompT gal* ($\lambda$ DE3) [*dcm*] Δ*hdsS* | (Studier & Moffattf, 1986) |
| **Plasmids** |  |  |
| pK18mob::sacB | *lacZa* Km<sup>r</sup> *sacB* *mob* | (Schäfer et al., 1994) |
| pK18mob2-EGFP | *lacZa* Km<sup>r</sup> *mob*, EGFP | (Schäper et al., 2016) |
| pSRKKm-EGFP | pSRKKm carrying EGFP coding sequence, Km<sup>r</sup> | (Schäper et al., 2016) |
| pWH844 | Protein expression vector containing N-terminal His-tag sequence Amp<sup>r</sup> | (Schirmer et al., 1997) |
| pPHU-EGFP | pPHU231 carrying EGFP coding sequence. Tc<sup>r</sup> | (Krol et al., 2016) |
| pLK114 | pPHU-EGFP with promoter region of *wgaA*, 298 bp UTR | (Charoenpanich et al., 2013) |
| pLK115 | pPHU-EGFP with promoter region of *wgeA*, 290 bp UTR | (Charoenpanich et al., 2013) |
| pPhaRdel | pK18mob::sacB carrying *phaR* flanking regions | This work |
| pPhbCdel | pK18mob::sacB carrying *phbC* flanking regions | This work |
| pPhaP1-EGFP | pK18mob2-EGFP carrying C-terminal portion of PhaP1 | This work |
| pPhaP2-EGFP | pK18mob2-EGFP carrying C-terminal portion of PhaP2 | This work |
| pPhaR-EGFP | pK18mob2-EGFP carrying C-terminal portion of PhaR | This work |
| pWH844-phaR | pWH844 carrying *phaR* coding sequence | This work |
| pPHU-PhaP1prCDS-mCherry | pPHU231 carrying complete *phaP1* gene, C-terminally fused to mCherry | This work |
| pBBRKM-EGFP | pSRKKm carrying EGFP coding sequence, EcoRI fragment containing *lacI<sup>q</sup>* and *lac* promoter removed | This work |
| pPHU-phaP1-EGFP | pPHU-EGFP with promoter region of *phaP1*, 192 bp UTR | This work |
| pPHU-phaP1-EGFP-proximal | pPHU-EGFP with promoter region of *phaP1*, 89 bp UTR | This work |
| pPHU-phaP2-EGFP | pPHU-EGFP with promoter region of *phaP2*, 176 bp UTR | This work |
| pPHU-phaP2-EGFP-proximal | pPHU-EGFP with promoter region of *phaP2*, 74 bp UTR | This work |
| pPHU-1961-EGFP | pPHU-EGFP with promoter region of *SMa1961*, 136 bp UTR | This work |
| pPHU-phaZ1-EGFP | pPHU-EGFP with promoter region of *phaZ*, 265 bp UTR | This work |
| pPHU-phaR-EGFP | pPHU-EGFP with promoter region of *phaR*, 135 bp UTR | This work |
| pPHU-guaB-EGFP | pPHU-EGFP with promoter region of *guaB*, 286 bp UTR | This work |
| pSRKKm-exoY-EGFP | pSRKKm-EGFP with promoter region of *exoY*, 473 bp UTR | This work |
| pSRKKm-exoL-EGFP | pSRKKm-EGFP with promoter region of *exoL*, 335 bp UTR | This work |
| pSRKKm-exoH-EGFP | pSRKKm-EGFP with promoter region of *exoL*, 297 bp UTR | This work |
| pSRKKm-ppdK-EGFP | pSRKKm-EGFP with promoter region of *ppdK*, 287 bp UTR | This work |
| pSRKKm-03815-EGFP | pSRKKm-EGFP with promoter region of *SMc03815*, 236 bp UTR | This work |
| pSRKKm-02240-EGFP | pSRKKm-EGFP with promoter region of *SMc02240*, 200 bp UTR | This work |
| pSRKKm-01834-EGFP | pSRKKm-EGFP with promoter region of *SMc01834*, 121 bp UTR | This work |
| pSRKKm-20609-EGFP | pSRKKm-EGFP with promoter region of *divK*, 270 bp UTR | This work |
| pSRKKm-panC-EGFP | pSRKKm-EGFP with promoter region of *panC*, 297 bp UTR | This work |
| pSRKKm-sdhC-EGFP | pSRKKm-EGFP with promoter region of *sdhC*, 198 bp UTR | This work |
| pSRKKm-smb21115-EGFP | pSRKKm with promoter region of *SMb21115*, 273 bp UTR | This work |
| pBBRKM-rhaD-EGFP | pBBRKM-EGFP with promoter region of *rhaD*, 287 bp UTR | This work |
| pBBRKM-zwf-EGFP | pBBRKM-EGFP with promoter region of *zwf*, 298 bp UTR | This work |
| pBBRKM-0319-EGFP | pBBRKM-EGFP with promoter region of *SMa0319*, 213 bp UTR | This work |
| pBBRKM-PacyP-EGFP | pBBRKM with promoter region of *acyP*, 285 bp UTR | This work |

**Table 2.**
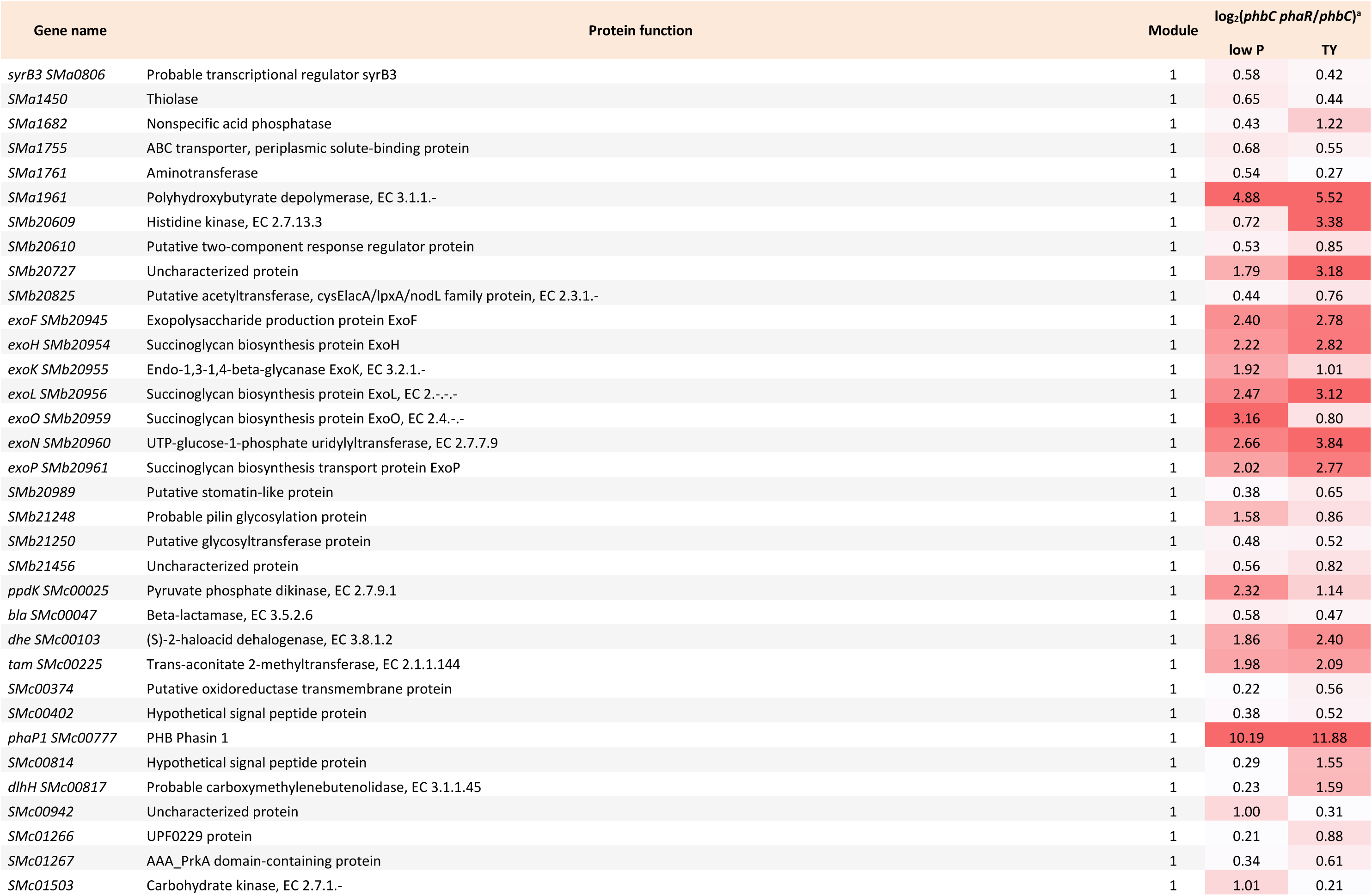
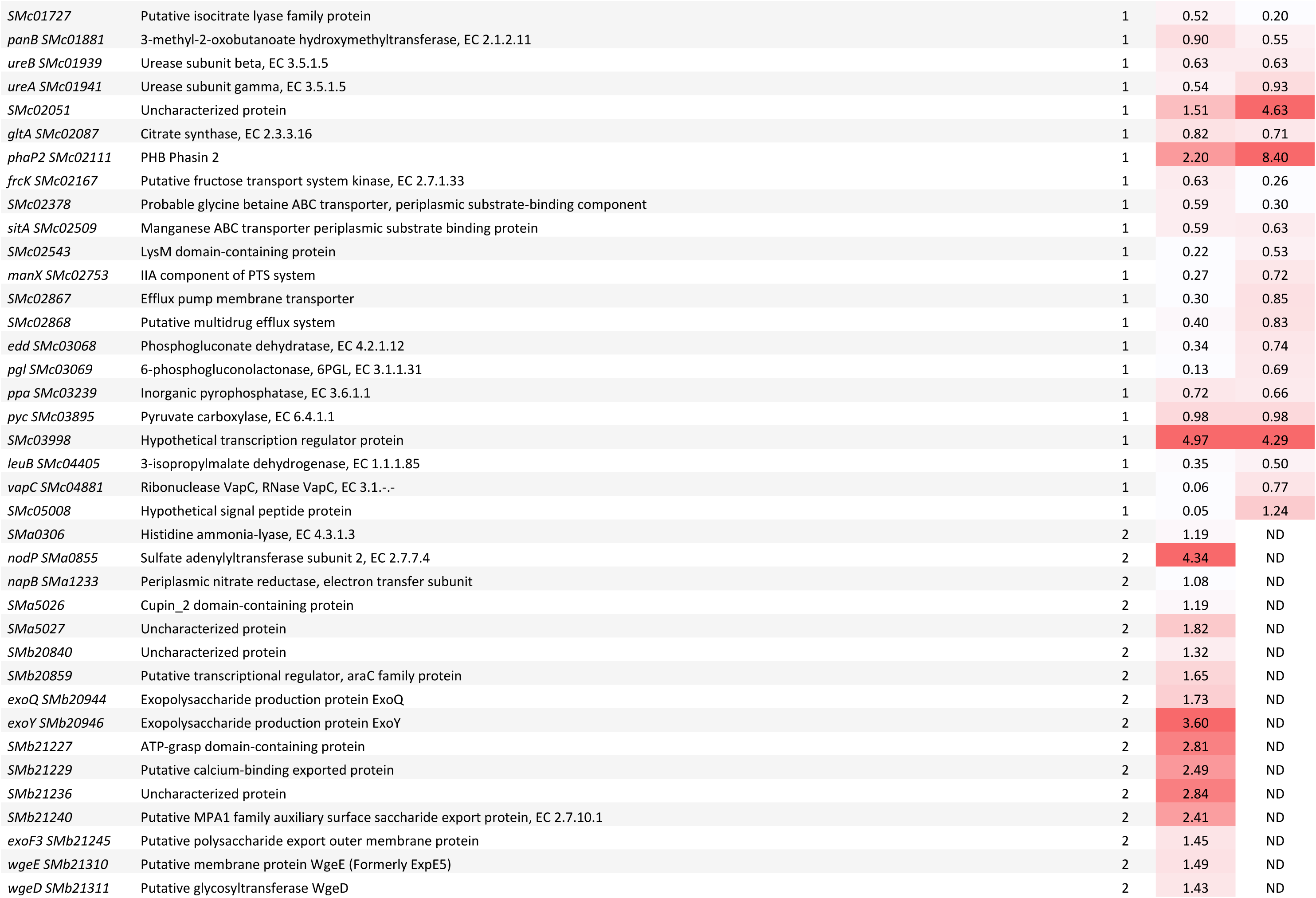
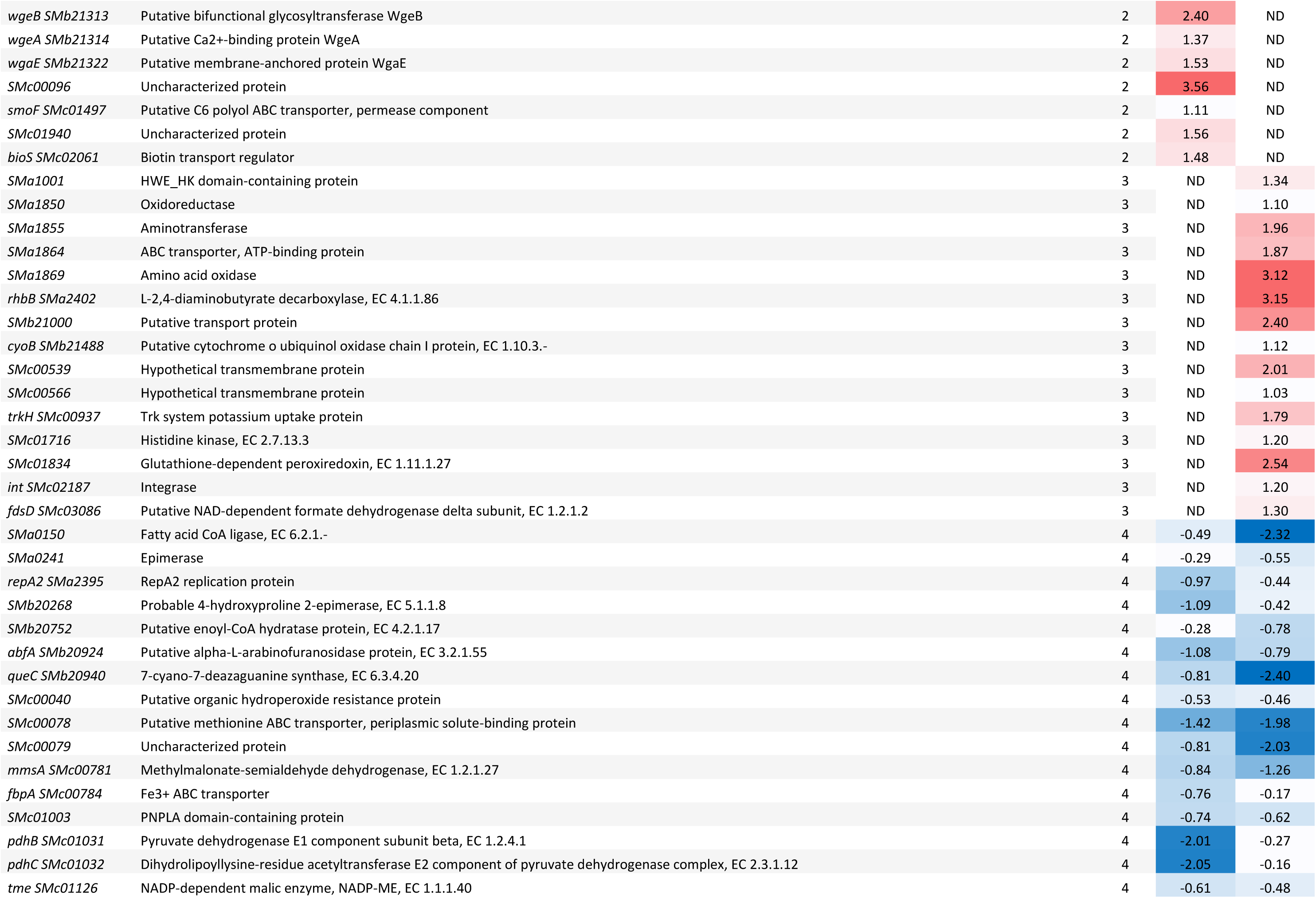
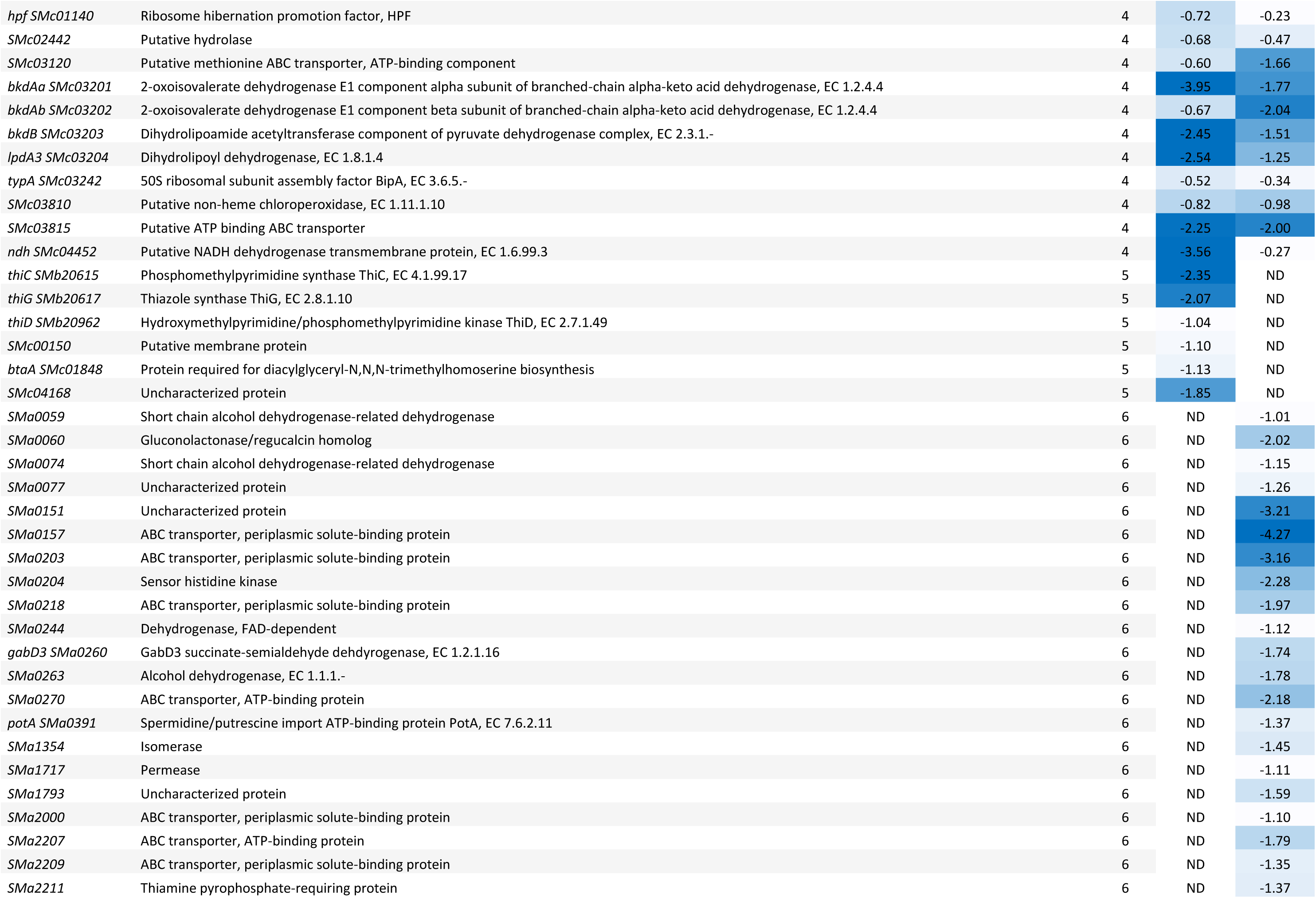
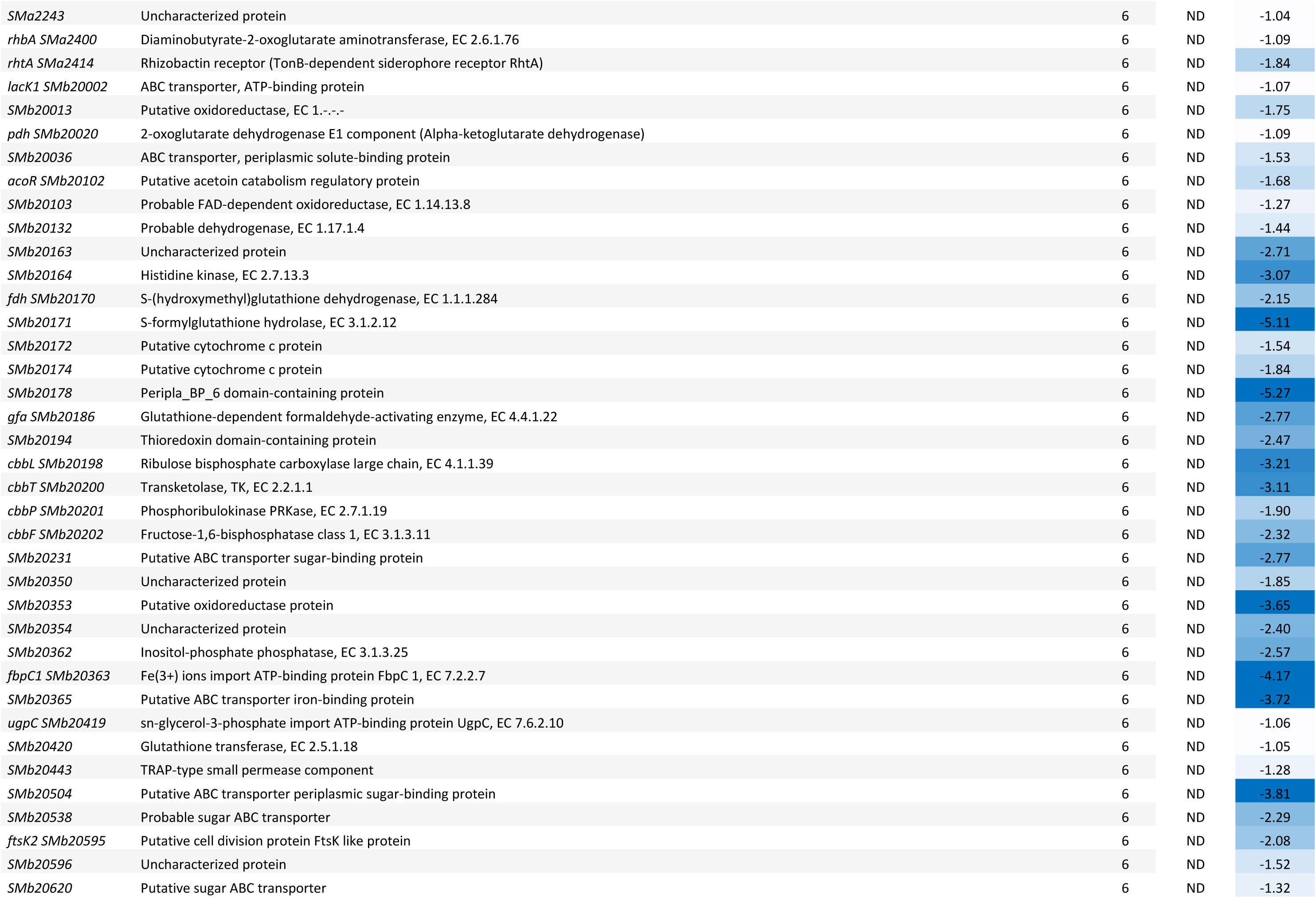
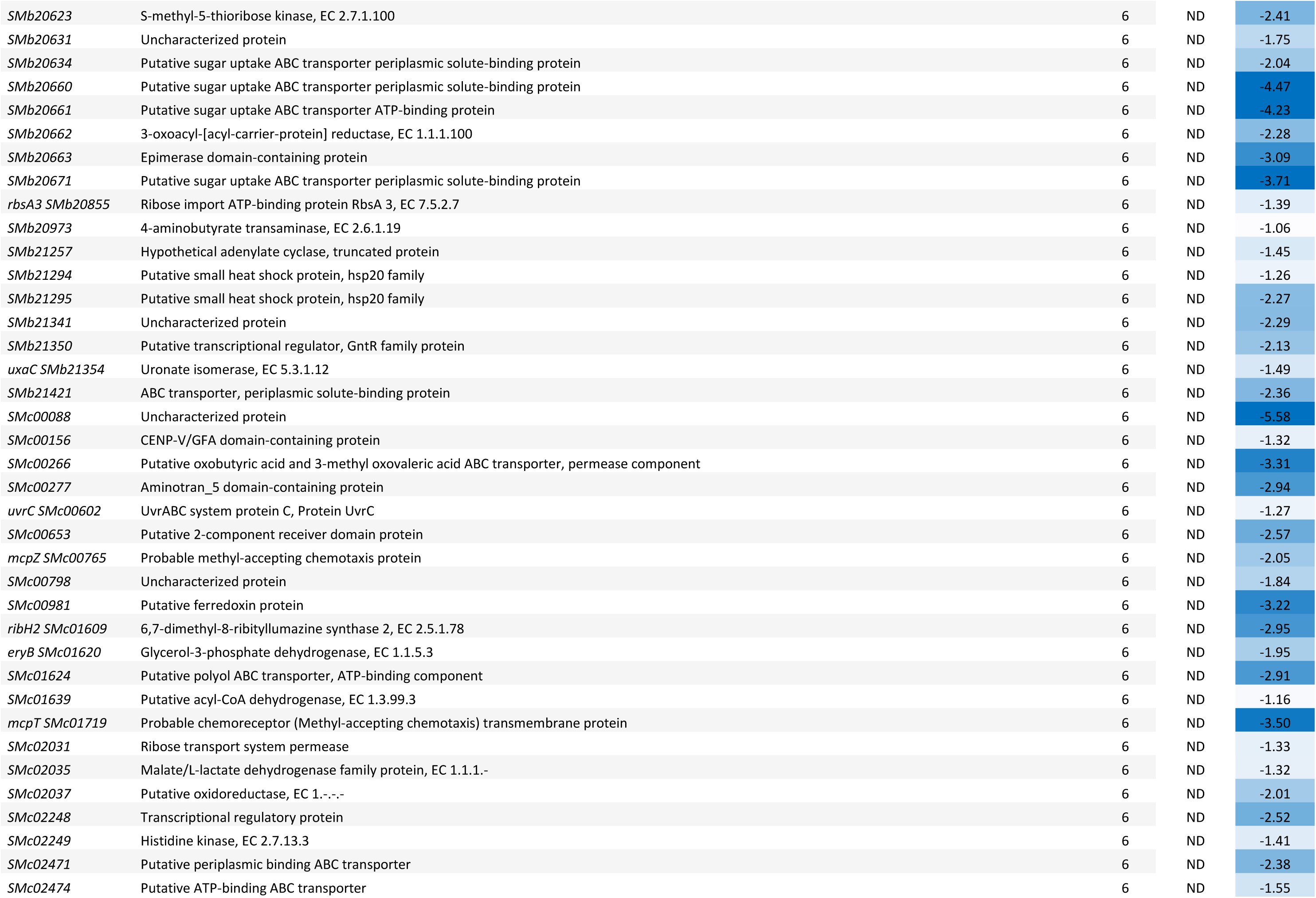
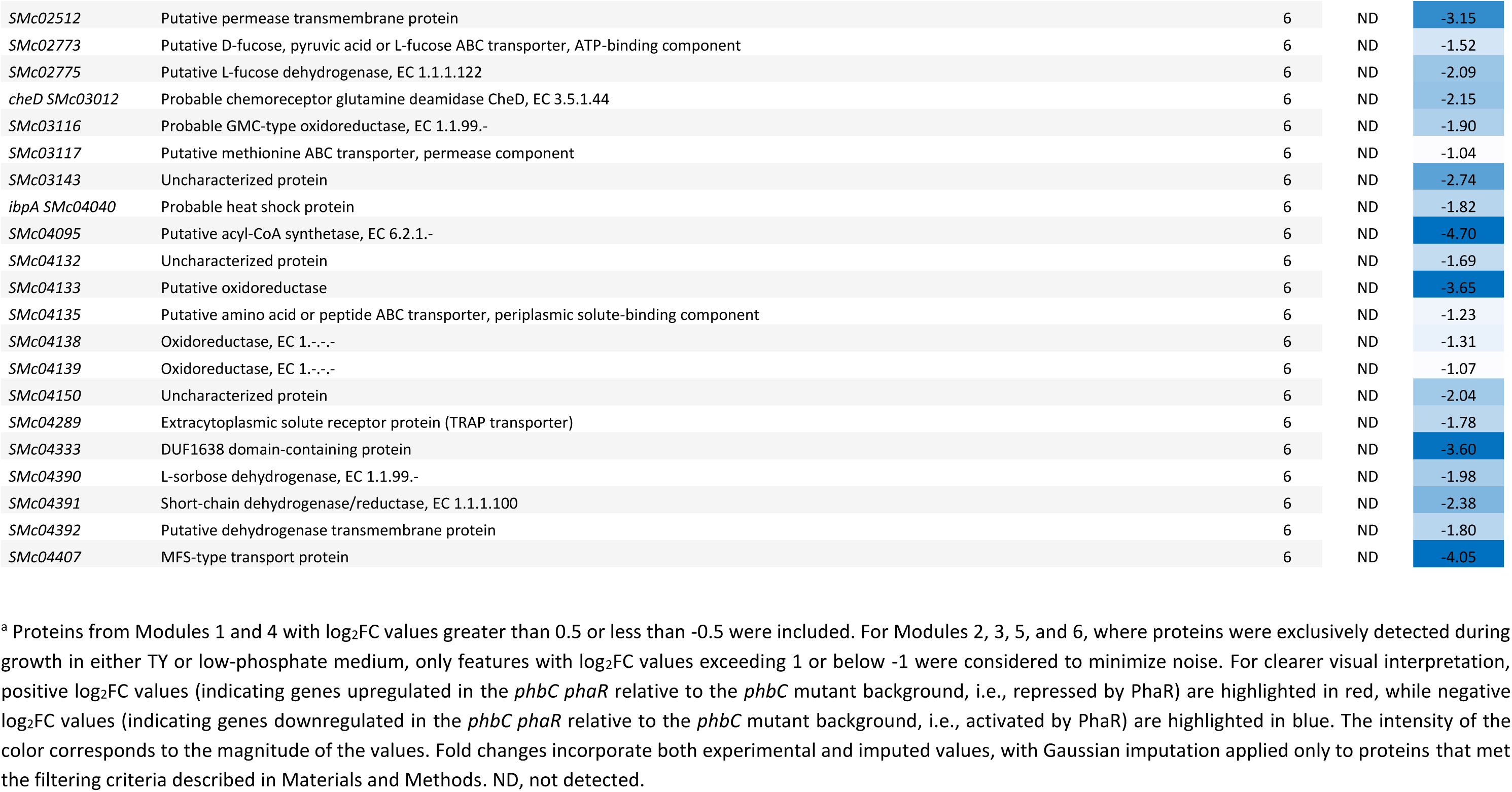
Fold changes of genes in Modules 1 to 6 in the *phbC phaR* compared to the *phbC* mutant under low-phosphate and TY media conditions.

Among proteins that clustered in Modules 4 to 6, components of the bacterial alpha-keto-acid dehydrogenase complexes, pyruvate dehydrogenase (encoded by the operon *pdh*) and beta-ketoacid dehydrogenase (encoded by the operon *bkd*), showed a strong positive regulation by PhaR (Table 2). While the structural components (EI- and EII proteins PdhA/B/C and BkdA/B) are specific for each complex, EIII proteins (LpdA proteins) are common components of all alpha-keto-acid dehydrogenases and likely interchangeable (Soto et al., 2001). Remarkably, ThiC/G/D, which are involved in the biosynthesis of thiamine pyrophosphate − a cofactor shared by the alpha-keto-acid dehydrogenase complexes − showed positive regulation by PhaR. However, these proteins were only detected in low P medium (Module 5), likely due to repression of their production by the exogeneous thiamine present in complex TY medium. Noteworthy, the levels of the Calvin–Benson–Bassham cycle-related proteins CbbF/P/T/L, detected only during growth in TY medium, were strongly positively influenced by PhaR. CbbA (whose coding gene in pSymB forms an operon together with *cbbFPTLSX*, *ppe* and *SMb20194*) did not cluster in Module 6 in the co-expression analysis, although it showed a significant upregulation upon *phaR* mutation in TY medium (supp Table 2). The methyl-accepting chemotaxis proteins McpT and McpZ (Zatakia et al., 2018) clustered in Module 6, which groups proteins detected exclusively in complex medium, with large fold-ratios indicating a strong activation by PhaR. Remarkably, 30% of the genes detected in TY medium as activated by PhaR (Module 6) are associated with membrane-transport functions (i.e., ABC-type transporter, TRAP transporter).

### Regulon-centric identification of the PhaR DNA binding motif in *S. meliloti* enables a genome-wide exploration of direct PhaR targets

The DNA-binding motif of PhaR homologs has been well characterized in several alpha-proteobacteria (Chou & Yang, 2010; Maehara et al., 2002; Nishihata et al., 2018). In *S. meliloti*, there is limited evidence suggesting that this motif is conserved. One study has shown that PhaR (AniA) acts as a repressor of the small non-coding RNA MmgR, which notably plays a role in regulating PHB accumulation (Lagares, Linne, et al., 2017). The promoter region of *mmgR* contains a putative binding site for PhaR that closely resembles previously published motifs (Ceizel Borella et al., 2018).

To identify the DNA binding motif recognized by PhaR in *S. meliloti*, we conducted a regulon-centric MEME search for conserved palindromic DNA sequences in the upstream regions of genes encoding proteins whose levels were affected by PhaR, specifically those included in Modules 1 to 6. We discovered the palindromic motif W_2_N_2_YGCRNYGCRN_2_W_2_, which was present in 49 promoter regions, with a clear enrichment in genes belonging to Module 1, and ∼30% of them (e.g., 22 genes out of 71 genes grouped in Module 1) having at least one copy in their upstream sequence (Fig. 4 and Table 3). Conversely, the motif was found in only 7 genes activated by PhaR (e.g., genes in modules 4 to 6). The high sequence identity shared between the identified motif and PhaR consensus binding sequences reported in related species (Fig. 4B) suggests that this sequence is likely the DNA sequence targeted by PhaR in *S. meliloti.* Our systemic approach to identifying the motif improved its definition and revealed extended AT-rich arms, which had not been reported so far in other species.

**Figure 4.**
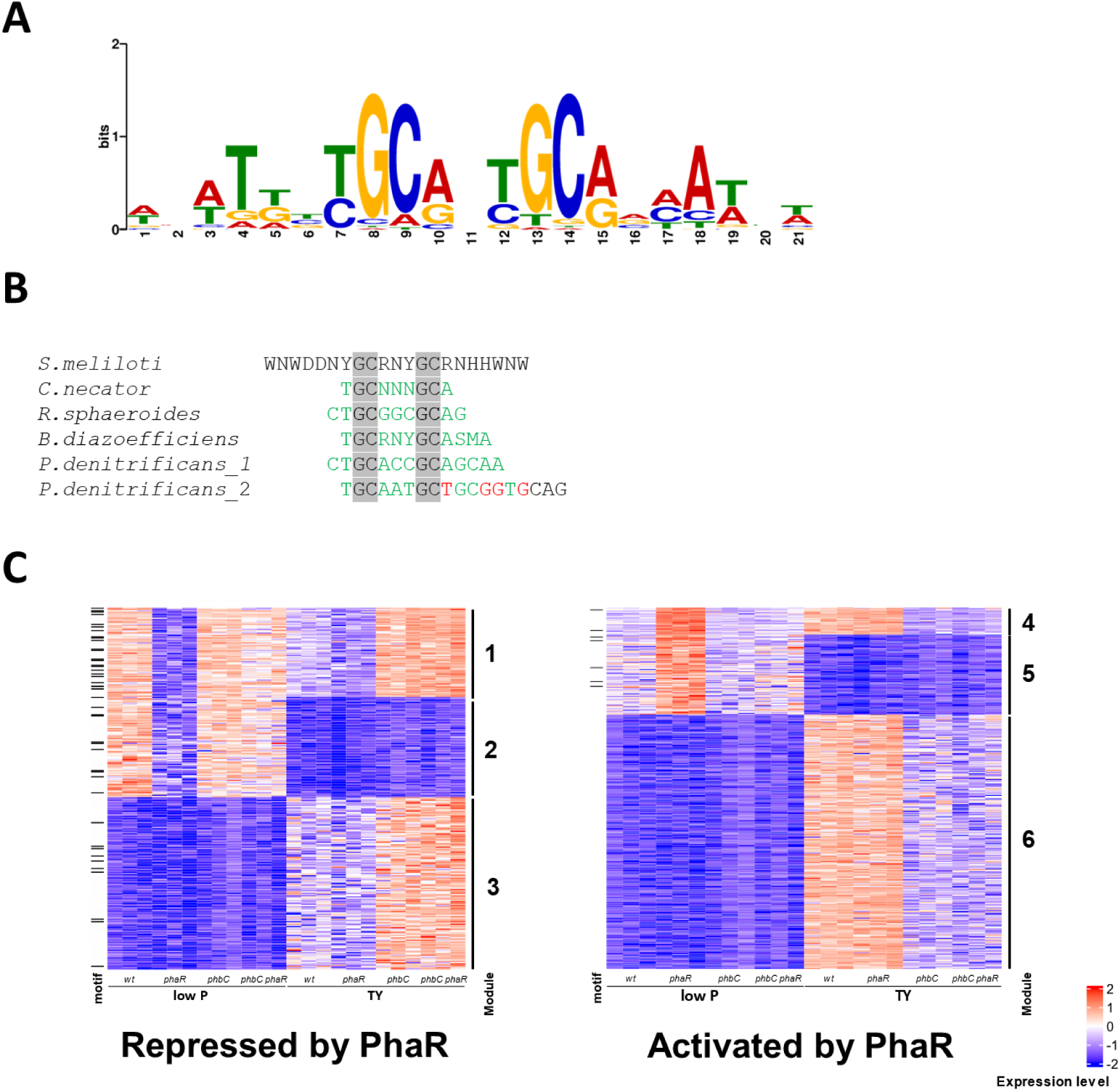
Regulon-centric identification of the PhaR target DNA binding motif in *S. meliloti.* **(A)** Consensus PhaR-binding site predicted using MEME (Bailey & Elkan, 1994) to search for conserved palindromic DNA sequences in the promoter region of genes that were clustered in Modules 1 to 6. **(B)** Alignment of PhaR-binding motifs experimentally characterized in multiple species. The conserved palindromic GC pairs are highlighted in gray. Positions in homologous motifs that match with the consensus sequence determined in *S. meliloti* are shown in green. Conversely, non-matching positions are shown in red. **(C)** Heatmaps depict the Z-scored abundances of PhaR-regulated proteins assigned to Modules 1 through 6 across strains grown in low P and TY media. Data from three biological replicates of each strain grown in each condition are displayed in separate adjacent columns. Features with at least one predicted PhaR DNA-binding motif in their promoter regions are indicated with horizontal lines on the left side of the plots. The Z-scored expression levels of each protein in each proteome (ranging from -2 to +2) are represented using the following color scale: blue indicates low expression, white represents intermediate expression, and red denotes high expression.

**Table 3.**
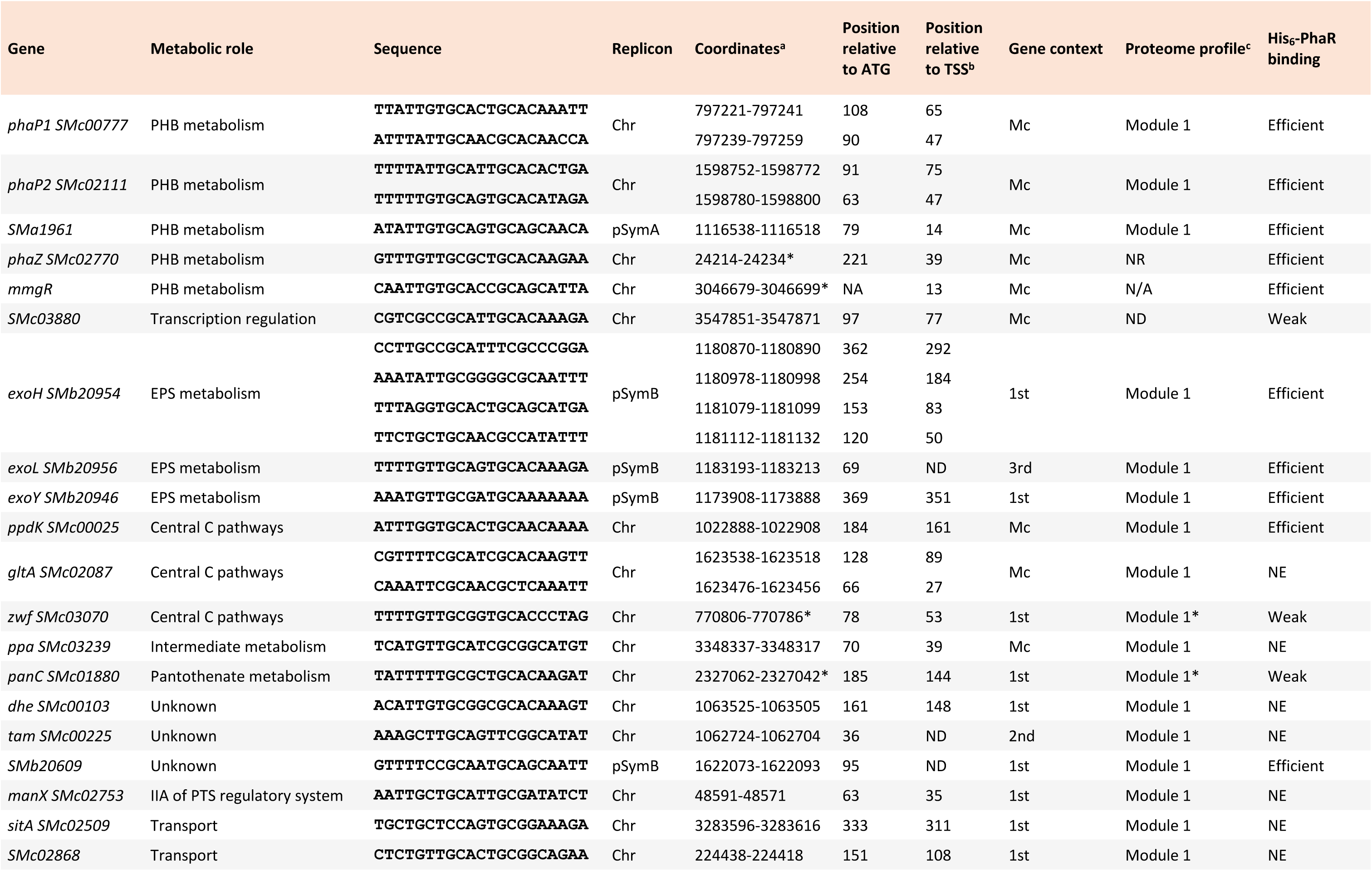
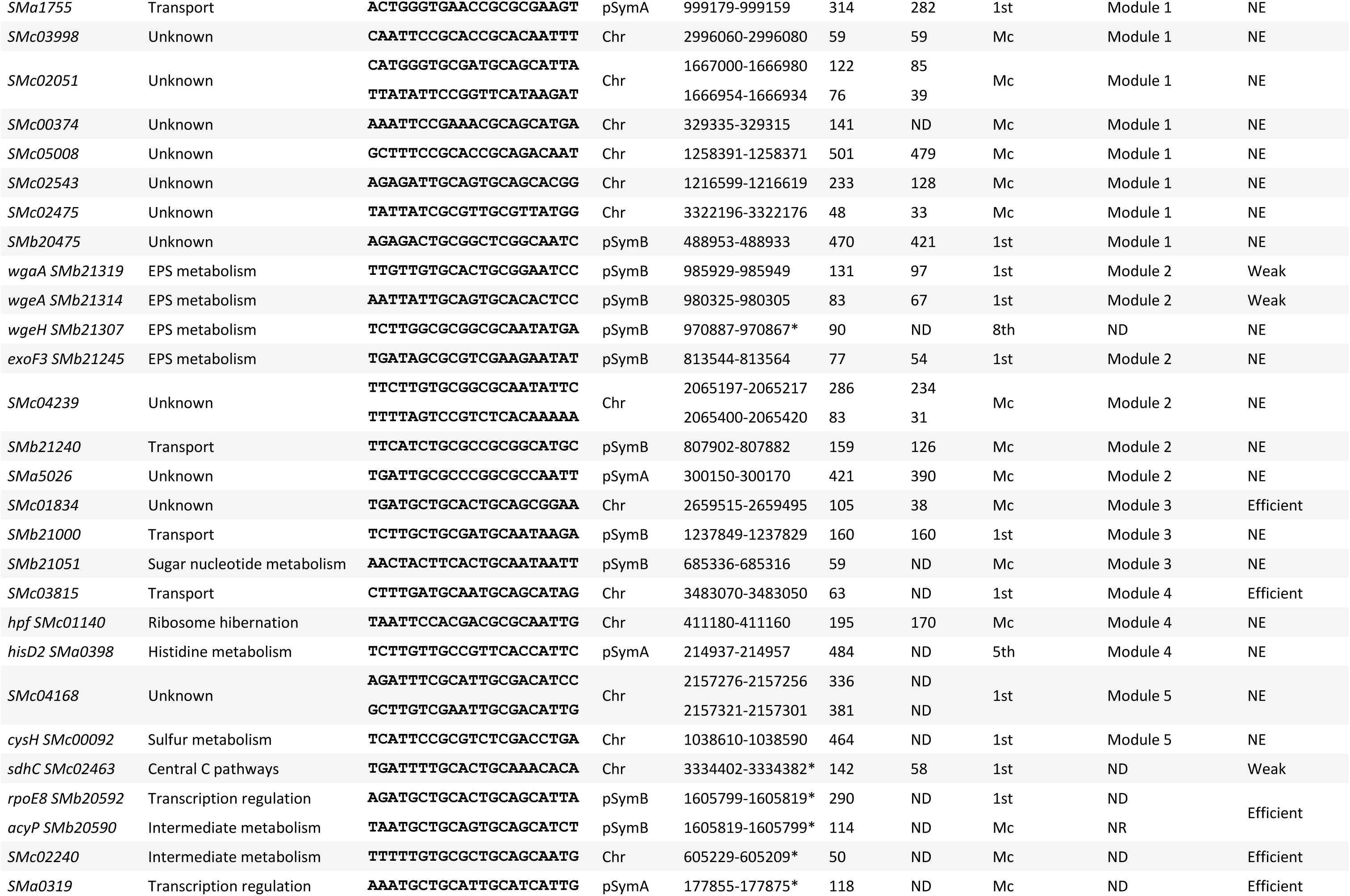
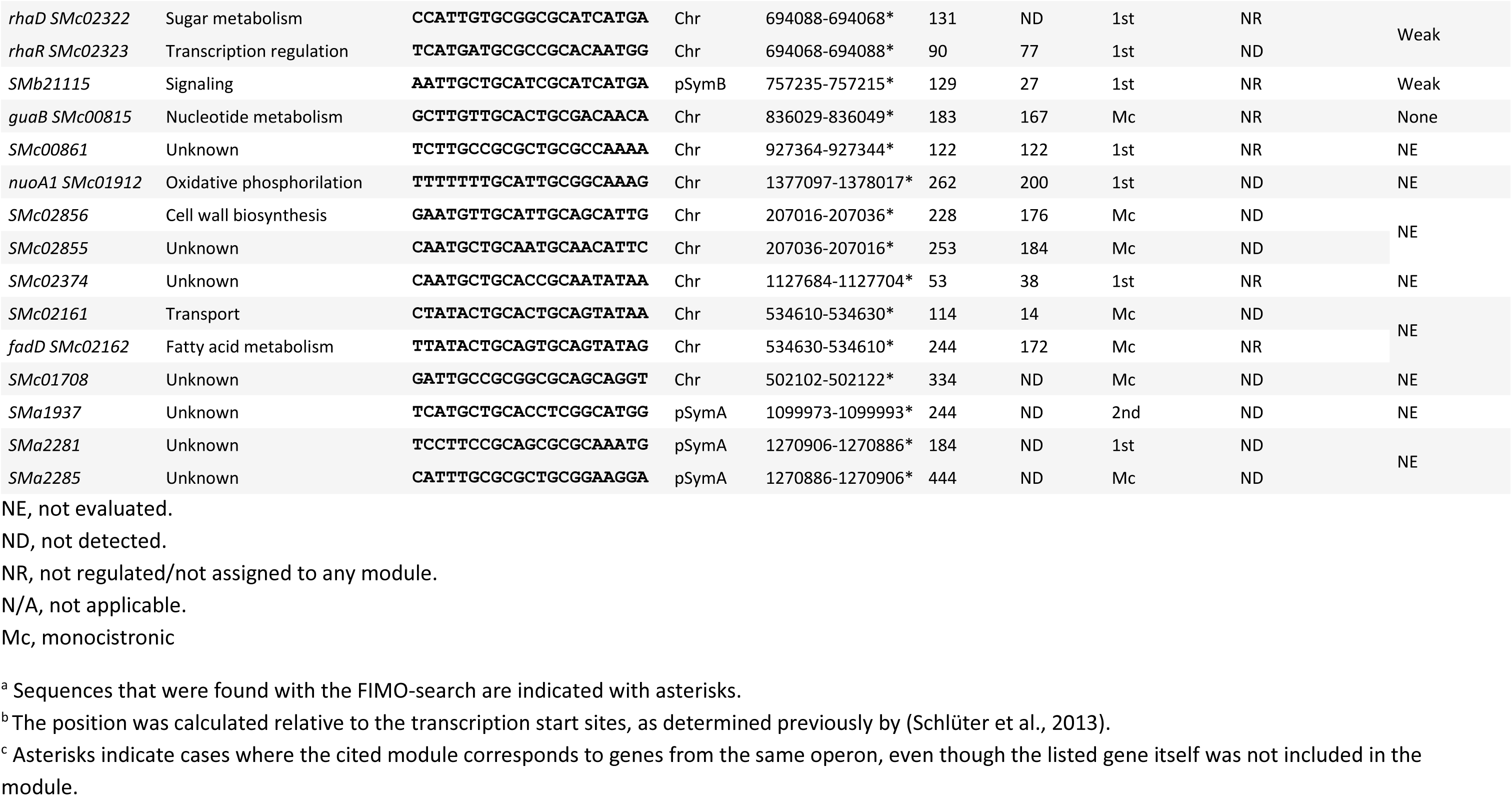
Promoter regions of PhaR-regulated genes containing PhaR-binding site-like sequence [TA]_2_N_2_[CT]GC[AG]N[CT]GC[AG]N_2_[TA]_2_.

Using FIMO, we then scanned the upstream regions of genes annotated in the *S. meliloti* genome for the conserved motif. This approach aimed to uncover additional PhaR target genes that may have remained undetected through our initial strategy, which relied on a limited set of proteins identified through proteomics. Since a motif search in upstream regions of genes putatively regulated by PhaR is likely to fail if it is not a monocistronic gene or the first gene of a polycistronic operon, this analysis sought to find the motifs in the promoter regions of such operons. Indeed, we identified a putative PhaR-binding motif upstream of the *zwf* gene (*SMc03070*), which encodes glucose-6P dehydrogenase and is the first gene in an operon followed by *pgl* and *edd* (whose encoded proteins clustered in Module 1). Similarly, *panC* (*SMc01880*) contains a putative PhaR-binding sequence and is positioned at the start of the panthotenate biosynthetic operon, which includes *panB* (Module 1). Table 3 lists the putative binding sites and their associated genes identified by the MEME search, as well as the 20 previously undetected binding sites with the highest score from the further FIMO search that were located in intergenic regions. Eleven of these sequences were further tested for binding to PhaR, as described below.

### PhaR senses PHB and regulates its metabolic modulon, targeting phasins and the degradative branch in *S. meliloti*

PHB-granule associated phasins PhaP1 and PhaP2, along with the PHB depolymerase SMa1961, displayed proteomic profiles across the tested mutant backgrounds and growth conditions that align with genes repressed by PhaR (Fig. 5A, top plots). These genes showed the strongest repression by PhaR in the proteomic analysis, with log_2_(fold change) ranging from 2 to 11 in the *phbC phaR* compared to the *phbC* mutant. The promoters of *phaP1*, *phaP2*, and *SMa1961* contain putative PhaR-binding motifs (Fig. 5B). Specifically, the *phaP1* and *phaP2* promoters each have two copies of the palindromic sequence; both located upstream of the -35 promoter element. In contrast, *SMc1961* has a single putative binding sequence that overlaps the -10 element. Furthermore, the genome-wide search using FIMO identified a putative binding site overlapping the -35 element in the sequence upstream of a second PHB depolymerase gene, *phaZ* (*SMc02770*) (Table 3, Fig. 5B). Although PhaZ did not exhibit changes at the proteomic level indicative of regulation by PhaR (Fig. 5A, top plots), we considered the possibility of an indirect effect that might mask this regulation and opted to investigate it further. Notably, our FIMO search also identified the presence of the motif in the promoter region of *mmgR* (Lagares, Linne, et al., 2017) (Fig. 5B). The activity of *phaP1* and *SMa1961* promoters, as assessed by low-copy number plasmid-borne promoter-*EGFP* fusions, correlated well with the corresponding protein abundances across the different mutant backgrounds, both under conditions that are permissive and non-permissive for PHB synthesis (with r^2^=0.9619 and r^2^=0.8092, respectively, Fig. 5A, bottom plots). Unlike PhaP1, however, PhaP2 was abundant only in PHB-producing, phosphate-limited wild type or *phaR* mutant cells, but not in cells of the *phbC* or *phbC phaR* strains grown in TY or low P media (Fig. 5A, top plots). This was in contrast to the high activation of the *phaP2* promoter in *phaR* and *phbC phaR* strains regardless of the growth medium (r^2^=0.6831, Fig. 5A, bottom plots). These results suggest that removal of PhaR − either through binding to PHB granules or through gene deletion − is sufficient to drive high levels of PhaP1 production. In contrast, the production and/or stability of PhaP2 appears to depend on both the absence of PhaR and presence of PHB granules, highlighting distinct post-transcriptional regulatory and/or stability mechanisms for each phasin. Noteworthy, the *phaZ* promoter activity displayed a profile pattern consistent with a gene typically repressed by PhaR, even though not correlating with PhaZ levels (r^2^=0.2429, Fig. 5A, bottom plots). This suggests that post-transcriptional mechanisms independent of PhaR operate on *phaZ* to finetune PhaZ abundance.

**Figure 5.**
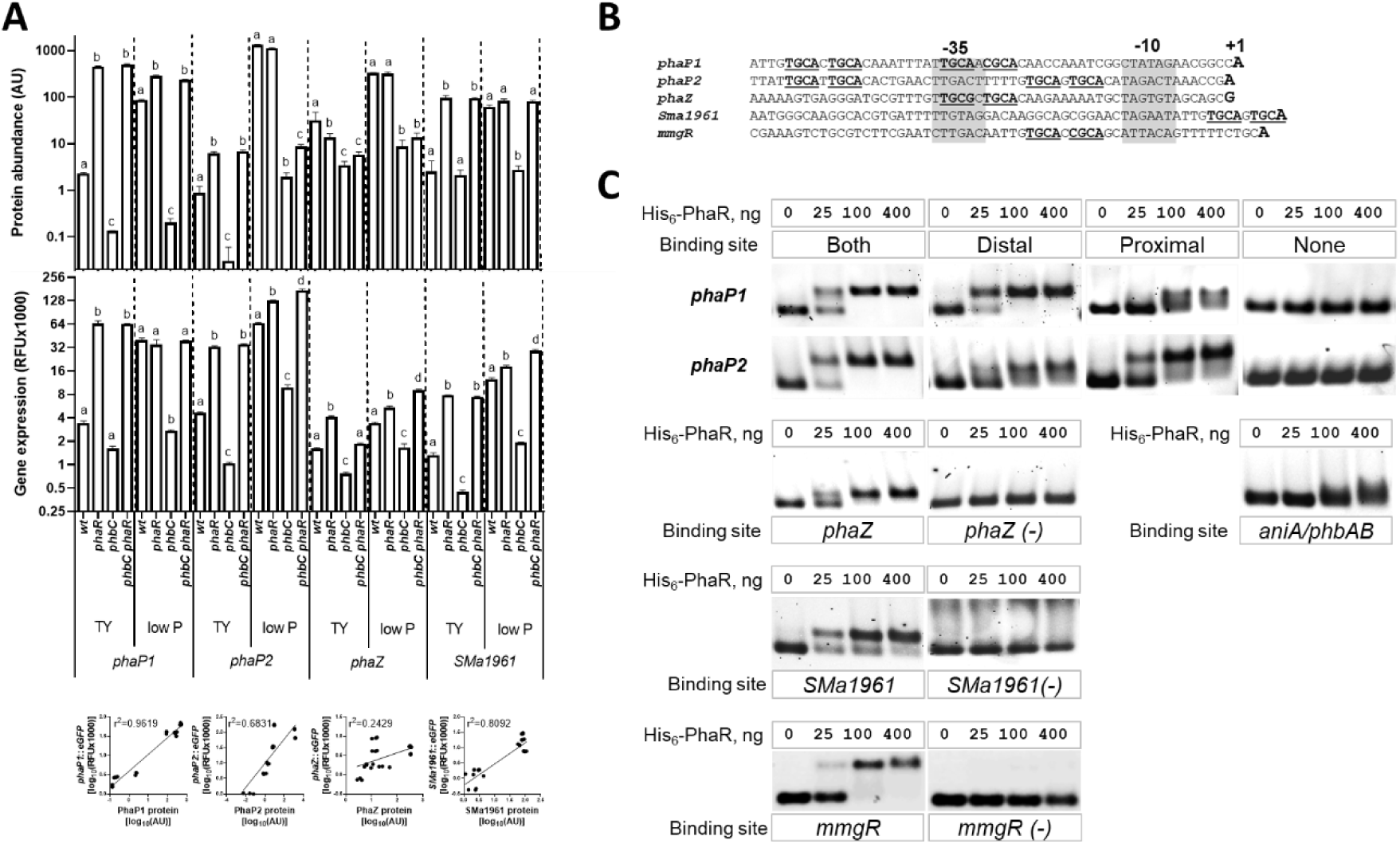
PhaR senses PHB and exerts direct control over its modulon. (A) (Top plots) Levels of PHB-associated phasins PhaP1, PhaP2, and PHB depolymerases PhaZ and SMa1961 across mutant backgrounds during growth in TY and low P media. Protein abundances are indicated in Arbitrary Units (AU), retrieved from proteomic data. Error bars represent standard deviation of three biological replicates. Statistical analysis was conducted using ANOVA followed by a permutation-based FDR correction for multiple-comparisons. Bars labeled with different letters indicate statistically significant differences (P-adj < 0.05). **(Central plots)** Promoter activity of phasins and PHB depolymerase genes. Fluorescence of promoter-*EGFP* fusions expressed from medium or low-copy number plasmid derivatives (see Table 1) was measured after 48 hs of cultivation. Error bars represent standard deviation of three to four biological replicates. Statistical analysis was conducted using ANOVA, followed by Tukey’s multiple-comparison test for each gene and growth condition. Bars labeled with different letters indicate statistically significant differences (P< 0.001). **(Bottom plots)** Correlation analysis between promoter activity of *phaP1*, *phaP2*, *phaZ* and *SMa1961* genes with levels of encoded proteins. The r^2^ values are indicated on the plots. **(B)** Promoter regions of phasins *phaP1/2*, PHB depolymerases *phaZ* and *SMa1961*, and *mmgR*. Transcription start site is shown with a letter in bold, - 35 and -10 promoter elements are marked by gray boxes, PhaR-binding palindromes are bolded and underlined. **(C)** EMSA assays with purified His_6_-PhaR and DNA fragments corresponding to wild type or modified sequences of promoters for *phaP1*, *phaP2*, *phaZ*, *SMa1961, mmgR* and *phaR/phbAB*.

Next, we tested the binding of His_6_-PhaR to the DNA fragments containing the promoter regions of *phaP1, phaP2, SMa1961*, *phaZ* and *mmgR* using an electrophoretic mobility shift assay (EMSA). PhaR binding was classified as efficient when a complete band shift occurred with 100 ng of His_6_-PhaR per reaction. Binding was considered low-efficiency if the shift was diffuse or required 400 ng of His_6_-PhaR. All promoter regions tested exhibited a clear shift when incubated with increasing amounts of PhaR (Fig. 5C). Excluding or introducing point mutations in the palindromic regions from each corresponding sequence abolished the shifts in the EMSA, providing strong evidence that these palindromes represent PhaR-binding sites. For both *phaP1* and *phaP2*, either the proximal or distal copy of the binding site was sufficient to promote PhaR binding in the EMSA.

Since autoregulation of *phaR* homologs has been reported in other bacteria (Maehara et al., 2002), we analyzed the *phaR/phbAB* intergenic region for binding by PhaR. Although two sequence motifs resembling the consensus site were found using FIMO, they fell below the threshold set for inclusion. Nonetheless, an electrophoretic mobility shift was observed, but only at the highest protein concentration tested (Fig. 5C). The promoter activity of *phaR* was slightly reduced in the *phaR* mutant background, whereas the promoter activity of *phbA* remained unaffected by *phaR* (data not shown). Additionally, no putative PhaR-binding site was detected in the promoter region of *phbC*, consistent with the failure to detect PhaR binding in the EMSAs and the unaltered promoter activity in response to PHB production or in case of *phaR* deletion. Taken together, these data suggest that PhaR regulates the expression of PHB depolymerase genes, but not the expression of the genes involved in PHB biosynthesis.

### PhaR links exopolysaccharide biosynthesis to PHB production

In *S. meliloti*, exopolysaccharide biosynthetic enzymes are strongly downregulated by PhaR, as revealed by the proteomic analysis (Figures 6A and 7A). Notably, regulation of the EPSII biosynthetic cluster at the proteome level is evident only during growth under low phosphate conditions, where the cluster becomes active due to the release of MucR-mediated repression (Bahlawane et al., 2008). The identification of a conserved PhaR-consensus binding sequence within the promoter regions of three genes associated to EPSI biosynthesis (e.g., *exoH*, *exoL* and *exoY*) and three genes related to EPSII production (e.g., *wgaA*, *wgeA* and *wgeH*) suggests direct regulation of these biosynthetic pathways by PhaR. To investigate this, we analyzed the binding of PhaR to the promoter regions of the EPSI biosynthesis genes *exoH*, *exoL* and *exoY*, and the galactoglucan genes *wgaA* and *wgeA* using EMSA.

**Figure 6.**
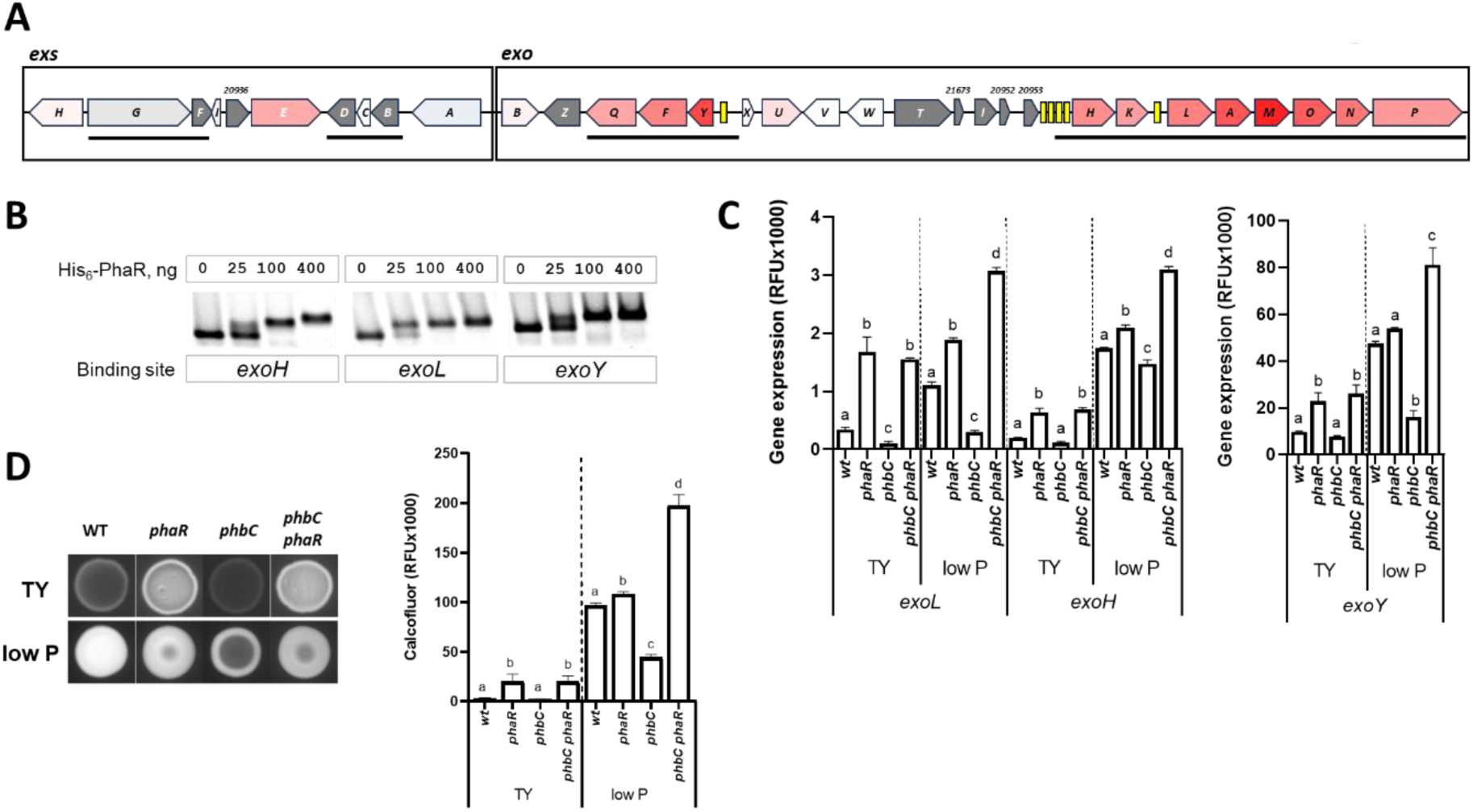
Succinoglycan (EPSI) biosynthesis regulation by PhaR. **(A)** Scaled genetic scheme showing proteome-level regulation of EPSI *exo* and *exs c*luster expression. Proteins encoded by genes highlighted in red showed significant upregulation in low P medium in the *phbC phaR* compared to the *phbC* mutant background, indicative of PhaR-mediated gene repression, with the intensity of the red color proportional to the log_2_(fold change) determined for each gene product. Proteins encoded by genes in gray were not detected. Operons, as described by Schlüter et al. (Schlüter et al., 2013), are indicated by solid lines below the scheme. Predicted PhaR-binding sites (listed in Table 3) are represented by yellow boxes. **(B)** EMSAs with purified His_6_-PhaR and the promoter regions of *exoH*, *exoL*, and *exoY* genes. **(C)** Promoter activity of EPSI biosynthesis genes. Fluorescence mediated by promoter-*EGFP* fusions was measured after 48 hs of cultivation. Error bars represent standard deviation of three to four biological replicates. Statistical analysis was conducted using ANOVA, followed by Tukey’s multiple-comparison test for each gene and growth condition. Bars labeled with different letters indicate statistically significant differences (P< 0.001). **(D)** EPSI production determined by calcofluor fluorescence quantification (right panel) of colonies that developed on agar plates of TY and low P media (left panel).

**Figure 7.**
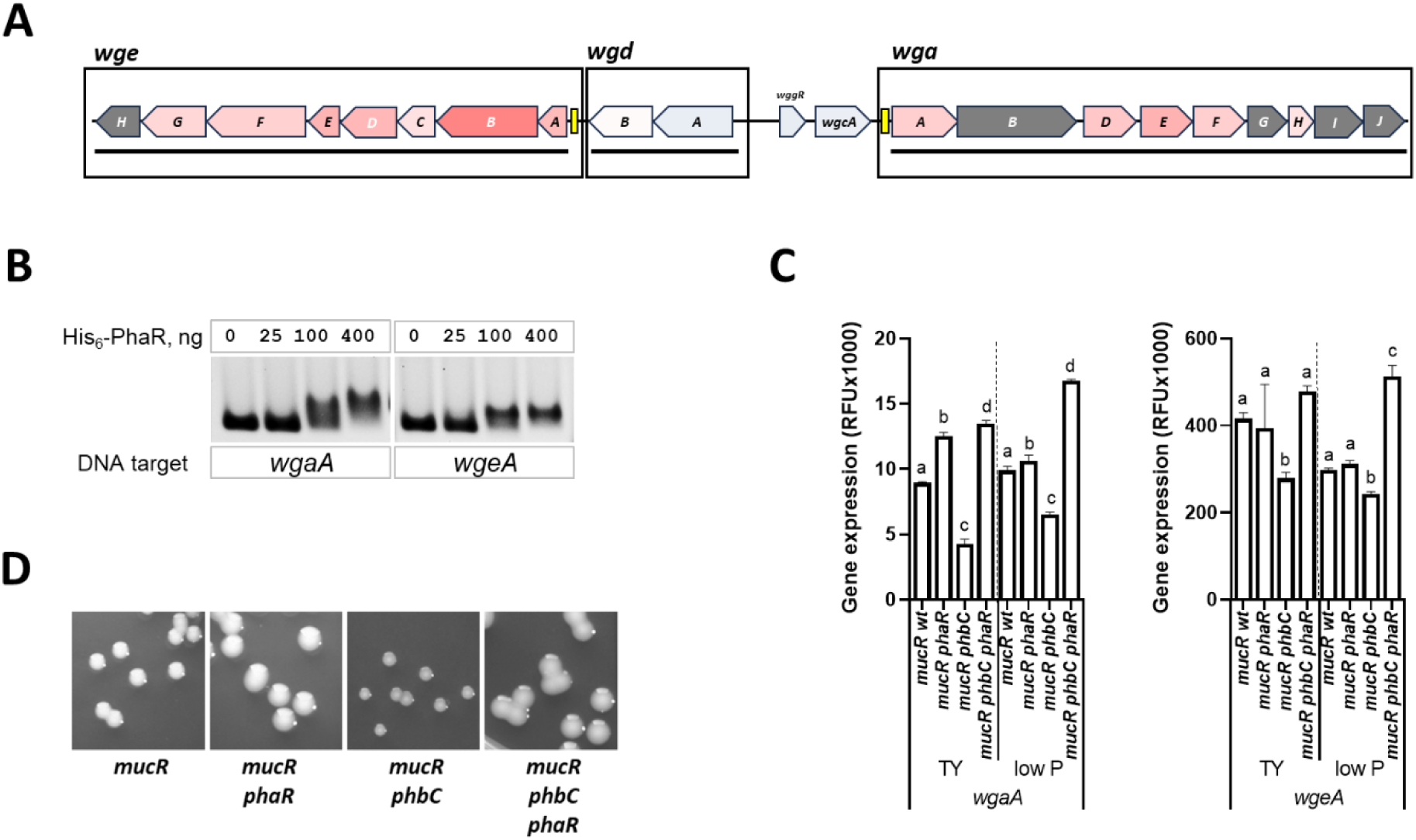
Galactoglucan (EPSII) biosynthesis regulation by PhaR. **(A)** Scaled genetic scheme showing proteome-level regulation of *wge*, *wgd* and *wga operons* of the EPSII biosynthesis gene cluster. Proteins encoded by genes highlighted in red showed significant upregulation in TY medium in the *phbC phaR* compared to the *phbC* mutant background, indicative of PhaR-mediated gene repression, with the intensity of the red color proportional to the log_2_(fold change) determined for each gene product. Proteins encoded by genes in gray were not detected. Operons, as described by Schlüter et al. (Schlüter et al., 2013), are indicated by solid lines below the scheme. Predicted PhaR-binding sites (listed in Table 3) are represented by yellow boxes. **(B)** EMSAs with purified His_6_-PhaR and the promoter regions of *wgaA* and *wgeA* genes. **(C)** Promoter activity of EPSII *wgaA* and *wgeA* biosynthesis genes. Fluorescence mediated by promoter-*EGFP* fusions was measured after 48 hs of cultivation. Error bars represent standard deviation of three to four biological replicates. Statistical analysis was conducted using ANOVA, followed by Tukey’s multiple-comparison test for each gene and growth condition. Bars labeled with different letters indicate statistically significant differences (P< 0.001). **(D)** Morphology of colonies developed by the indicated strains grown on TY agar.

The promoter regions of EPSI biosynthesis genes *exoH*, *exoL* and *exoY* efficiently bound PhaR (Fig. 6B). Thus, regulation of these promoters by PhaR likely controls the expression of the *exoHKLAMNOP* and *exoYFQZ* operons, which include 12 out of the 19 genes involved in EPSI biosynthesis. Promoter-*EGFP* assays further showed increased transcription of *exoY, exoL* and *exoH* in *phaR*-deficient strains (Fig. 6C). During growth in TY medium, these promoters were repressed in presence of *phaR,* with strongest repression observed in the *phbC* mutant. To analyze EPSI production, we performed growth assays in presence of Calcofluor, a fluorescent dye that binds to this EPS and allows its production to be assessed by the fluorescence of agar cultures (Fig. 6D). During growth on TY medium, the *phaR* strain exhibited significantly higher fluorescence than the wild type, suggesting enhanced EPSI production in the absence of PhaR. In low P medium, both the wild type and *phaR* strains displayed similar fluorescence, suggesting comparable levels of EPSI production. Thus, PhaR reduces EPSI production by repressing the transcription of the *exoYFQZ* and *exoHKLAMNOP* operons in PHB-dependent manner.

Furthermore, we observed weak PhaR binding at the promoter region of *wgaA* (Table 3, Fig. 7B). The *wgaA* promoter drives the expression of 10 *wga* genes involved in the biosynthesis of EPSII (Becker et al., 1997). In Sm2011 and its derivatives, EPSII biosynthesis genes, including *wgaA*, are repressed by the transcriptional regulator MucR (Bahlawane et al., 2008). To analyze the regulation of EPSII biosynthesis by PhaR and PHB, *phaR* and *phbC* mutations were introduced into a *mucR* deletion strain. Deletion of *phaR* in the *mucR* strain resulted in a slight increase in *wgaA* promoter activity; however, deletion of *phbC* reduced the P*_wgaA_*-*EGFP* fluorescence signal by approximately two-fold (Fig. 7C). In contrast, the *wgeA* promoter, which drives expression of the *wge* operon, did not display *phaR*-medited regulation in the *mucR* strain. Although the regulation of *wgaA* was moderate, the mucoidity of the *mucR* strain increased notably upon deletion of *phaR* and decreased upon mutation of *phbC* (Fig. 7D).

Remarkably, *ppa*, encoding the inorganic pyrophosphatase − an enzyme catalyzing the hydrolysis of pyrophosphate (PPi) to phosphate − was associated with Module 1. This reaction is essential for maintaining low PPi levels, a condition necessary to thermodynamically favor the activation of phosphate-sugars to nucleotide sugars (e.g., ExoN-catalyzed activation of glucose-1P to UDP-glucose, a precursor of EPS biosynthesis). Notably, an PhaR-binding motif was identified in the promoter region of *ppa*, suggesting that *ppa* is a direct target repressed by PhaR.

### The PhaR regulon extends beyond PHB and EPS metabolism

Putative consensus binding sites for PhaR were found in the promoter regions of the Module 1-associated genes *ppdK*, *gltA* and *manX* (Table 3), suggesting direct regulation by PhaR. The binding of PhaR to the *ppdK* promoter was tested as a proxy to confirm a direct involvement of PhaR in these processes. Efficient binding was detected between PhaR and the promoter region of *ppdK* (Fig. 8A), and *ppdK* promoter driven expression of the *EGPF* reporter was dependent on *phaR* and correlated well with PpdK levels in both media (r^2^=0.9393, Fig. 8B, bottom plots).

**Figure 8.**
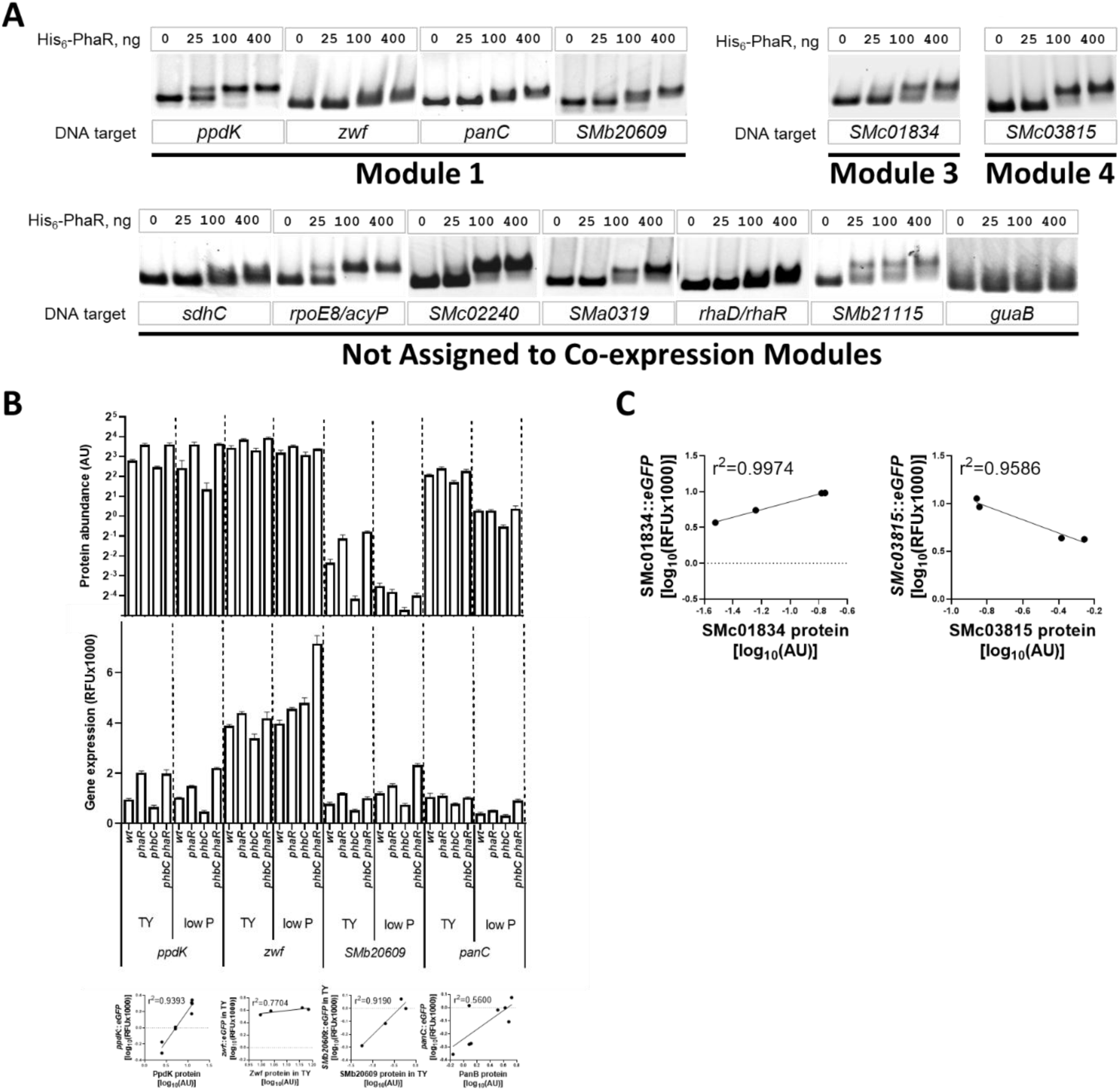
The PhaR regulon extends beyond PHB and EPS metabolism. **(A)** EMSA assays with purified His_6_-PhaR and DNA fragments corresponding to sequences of promoters for *ppdK*, *zwf*, *panC*, *SMb20609, SMc01834, SMc03815, sdhC, SMc02240, SMa0319, rhaD/R, SMb21115 and guaB.* **(B) (Top plots)** Levels of Module 1-clustered proteins PpdK, Zwf, Smb20609 and PanB across mutant backgrounds during growth in TY and low P media. Protein abundances are indicated in Arbitrary Units (AU), retrieved from proteomic data. Error bars represent standard deviation of three biological replicates. Statistical analysis was conducted using ANOVA followed by a permutation-based FDR correction for multiple-comparisons. Bars labeled with different letters indicate statistically significant differences (P-adj < 0.05). **(Central plots)** Promoter activity of the corresponding genes. Fluorescence of promoter-*EGFP* fusions expressed from medium or low-copy number plasmid derivatives (see Table 1) was measured after 48 hs of cultivation. Error bars represent standard deviation of three to four biological replicates. Statistical analysis was conducted using ANOVA, followed by Tukey’s multiple-comparison test for each gene and growth condition. Bars labeled with different letters indicate statistically significant differences (P< 0.001). **(Bottom plots)** Correlation analysis between promoter activity of *ppdK*, *zwf*, *SMb20609* and *panC* genes with levels of encoded proteins (except for *panC*, where correlation was calculated with PanB levels). The r^2^ values are indicated on the plots. **(C)** Correlation analysis between promoter activity of *SMc01834* and *SMc03815* genes with levels of encoded proteins. The r^2^ values are indicated on the plots.

PhaR was also found to exert mild repression on levels of Zwf, Pgl and Edd enzymes catalyzing the initial steps of sugar catabolism through the Entner-Doudoroff (ED) pathway, which results in glyceraldehyde-3P and pyruvate as end products (Figure 8B, Table 2). *zwf*, *pgl* and *edd* are organized in an operon and a putative binding site for PhaR was found immediately upstream to the -35 box of the leading promoter (Table 3). Weak binding of His_6_-PhaR to the *zwf* promoter region was confirmed through an EMSA (Fig. 8A), and regulation at the transcriptional level was supported by changes in fluorescence mediated by the P*_zwf_*-*EGFP* reporter fusion in TY medium but not in low P (r^2^=0.7704, Fig. 8B, bottom plots). While Zwf was not clustered along with Pgl and Edd in Module 1 (e.g., displaying a regulatory pattern compatible with PhaR repression subjected to PHB modulation) its levels do increase in the *phbC phaR* compared to the *phbC* mutant background (ANOVA, p-adj<0.05). Though the fold changes for Zwf, Pgl, and Edd were rather modest, even subtle adjustments in enzyme levels may contribute to the overall metabolic response governed by PhaR.

From Module 1, SMb20609 showed a strong dependence on *phaR*. Efficient binding of His_6_-PhaR to the *SMb20609* promoter region (Table 3) demonstrated by an EMSA (Fig. 8A), as well as the activity pattern of a P*_SMb20609_-EGFP* reporter fusion in wild type, *phbC phaR* and *phbC* strains during growth in TY and low P media strongly suggest direct regulation of *SMb20609* by PhaR at the transcriptional level (r^2^=0.9190, Fig. 8B, bottom plots).

As noted above, the FIMO search identified an PhaR-binding motif within the *panC* promoter region (Table 3). *panC* is the first gene in an operon that also includes *panB* coding for 3-methyl-2-oxobutanoate hydroxymethyltransferase, which like PanC is associated with Module 1 (Table 2). This enzyme catalyzes the initial step in the biosynthesis of pantothenate − a precursor of Coenzyme A − after diverging from the valine biosynthesis pathway. An EMSA revealed a weak interaction between PhaR and this motif (Fig. 8A) and P*_panC_*-*EGFP* reporter fusion activity suggests direct transcriptional regulation of this operon by PhaR), in good correlation with the proteomic pattern observed for PanB (r^2^=0.56, Fig. 8B, bottom plots).

The promoter regions of *SMc01834* and *SMc03815* were selected as representatives to analyze the direct binding and regulation by PhaR to promoters from genes that contain a PhaR-consensus binding motif and belong to Modules 3 and 4, respectively. Both promoters bound His_6_-PhaR efficiently (Fig. 8A). While the P*_SMc01834_-EGFP* fusion reported promoter activites in all mutant backgrounds that correlated well with the corresponding protein abundances in TY (r^2^=0.9974, Fig. 8C, bottom plots), the activities of the P*_SMc03815_-EGFP* fusion followed a pattern consistent with a PhaR-repressed gene. This contrasts with the behaviour observed for SMc03815 in the proteome analysis and is reflected by a negative correlation index (r^2^=0.9586, slope = -0.6945, Fig. 8C, bottom plots).

Finally, we explored the potential binding of PhaR to promoters where a consensus binding motif was detected with FIMO, specifically in genes encoding proteins that were either absent in the proteome or did not cluster in co-expression modules. EMSAs demonstrated that His_6_-PhaR bound to P*_sdhC_*, P*_SMc02240_*, P*_SMa0319_*, P*_SMb21115_*, and to the intergenic regions between *rpoE* and *acyP*, as well as *rhaD* and *rhaR*, with varying efficiency (Fig. 8A). However, no binding was detected between His_6_-PhaR and P*_guaB_* (Fig. 8A).

## Discussion

### PhaR’s dual role as a PHB sensor and gene regulator is likely an ancient, conserved trait

The role of PhaR in carbon flow partitioning in *Sinorhizobium meliloti* was first proposed by Povolo & Casella (Povolo & Casella, 2000a). Subsequent studies on PhaR homologs in proteobacteria, including *P. denitrificans* and *Cupriavidus necator*, revealed its function in coordinating PHB accumulation with metabolism by acting as both a transcriptional regulator and PHB sensor (Maehara et al., 2002; Pötter et al., 2002).

Despite these insights, a comprehensive analysis of the PhaR regulon remains lacking. Here, we investigated PhaR’s role in PHB sensing and gene regulation in *S. meliloti*. Using co-expression network analysis and proteomics across mutant backgrounds, we uncovered a PHB-dependent PhaR-mediated global response. Our approach improved sensitivity over traditional differential expression analyses (Horvath, 2011). Notably, 30-35% of identified proteins in low P and TY media showed *phaR*-dependent production (ANOVA, P-adj<0.05). We confirmed PhaR’s interaction with PHB through in vitro biochemical assays and in vivo co-localization, supporting a model where PhaR switches between active (free) and inactive (PHB-bound) states. Co-expression network analysis identified ten gene modules, with six displaying *phaR*-dependent regulation in a PHB-dependent manner. As expected, intermediate metabolism proteins emerged as key regulatory targets. Our findings highlight a pivotal role for PhaR as a central hub in *S. meliloti* which -through sensing of PHB levels-regulates central C-pathways and EPS production, both in connection with other cellular processes.

### Overlap between the PhaR regulon and PHB modulon varies across species

*S. meliloti* PhaR was found to repress the transcription of phasins and PHB depolymerase genes by binding in the vicinity of predicted -35 and -10 promoter elements, similar to PhaR homologs from Gram-negative bacteria studied so far (Maehara et al., 2002; Pötter et al., 2005). The direct regulatory link between PHB and phasin expression probably arose more than once during evolution, since it also operates in PHB producing *Bacillus* species, where phasin gene regulation function of PhaR is fulfilled by a non-homologous PHB- and DNA-binding protein PhaQ (Lee et al., 2004). Presence of multiple phasins is a widespread feature of PHB-producing bacteria. For example, *C. necator* and *B. diazoefficiens* possess seven and four phasins, respectively, which are expressed at different levels (Pfeiffer & Jendrossek, 2011, 2012; Pötter et al., 2004; Quelas et al., 2016). Such redundancy infers a complex control of granule proteome respective to PHB accumulation and growth conditions. In *Cupriavidus*, probably not all phasin genes are regulated by *phaR*, since PhaR was shown to bind in the promoter regions of *phaP1* and *phaP3*, but not *phaP2* and *phaP4* (Pötter et al., 2005). In *S. meliloti*, PhaR regulated both *phaP1* and *phaP2*. Although *S. meliloti* phasins differ in size (148 and 114 amino acid residues for PhaP1 and PhaP2, respectively) and share only 28% sequence identity, their structures are highly conserved and they share a common function in PHB binding (C. Wang, Sheng, et al., 2007). Repression by PhaR was identified as the main regulatory mechanism for *phaP1*. However, expression of *phaP2* was also modulated by other unknown factors linked to PHB-production, likely the small RNA MmgR (Lagares, Linne, et al., 2017). The observed differences in regulation suggest a degree of functional divergence among *S. meliloti* phasins.

Unlike in *B. diazoefficiens*, in *S. meliloti* PhaR targets PHB degradation genes but does not seem to directly regulate the activity of PHB biosynthetic enzymes (Quelas et al., 2016, 2024). The significant decrease in stored PHB upon *phaR* mutation under PHB-permissive conditions in low P medium (Fig. 1B), suggests incomplete PhaR sequestration by PHB. Under these conditions, residual regulatory activity of a small proportion of DNA-bound PhaR molecules may be critical for properly adjusting cellular PHB-depolymerase activity, ensuring optimal PHB accumulation levels.

Negative autoregulation by PhaR in *P. denitrificans* and *C. necator* was suggested based on the protein’s ability to bind in the promoter region of *phaR*, although the expression levels in the presence or the absence of *phaR* were not measured (Maehara et al., 2002, Pötter et al., 2002). A qRT-PCR analysis in C*. necator* showed that the transcript abundance of *phaP*, but not *phaR*, was dependent on PHB production, making it unlikely that *phaR* is regulated in the same manner as phasin genes (Lawrence et al., 2005). In *R. sphaeroides*, negative autoregulation of *phaR* was directly observed, as the mutation in the gene increased its own transcription to about 130 % of wildtype levels (Chou et al., 2009). In our study, the promoter activity of *phaR* was analyzed in both *phaR*^+^ and *phaR* strains, and the results suggested a weak positive autoregulation. Binding of PhaR to its own promoter region in EMSA assays was weak, similar to the situation in *P. denitrificans*, where binding of PhaR to the *phaR* promoter was reported to be approximately 10 times less efficient than binding to the *phaP* promoter region (Maehara et al., 2002). Thus, the strength of regulation by PhaR is likely to differ between phasin gene promoters and their own promoters, suggesting that negative autoregulation of *phaR* homologs is probably not a conserved feature among PHB-producing bacteria.

### PhaR-mediated crosstalk between EPS and PHB production

Dramatically reduced succinoglycan production was reported previously for *S. meliloti phbC* strains, unable to produce PHB (C. Wang, Saldanha, et al., 2007). This effect was also observed in the Calcofluor assay. Several explanations for this phenomenon have been proposed, including regulation via the ExoS-ChvI two-component system in response to increased levels of acetyl phosphate (Trainer et al., 2010) or the coupled regulation of EPSI and PHB biosynthesis through an unknown mechanism (Povolo & Casella, 2009b). In this study, we clearly identified *phaR* as the factor responsible for this effect, as inactivation of *phaR* in *phbC* strain restored EPSI production (Fig. 5D). His_6_-PhaR was able to bind to the promoter regions of *exo* genes, and PhaR repressed their transcription *in vivo*. Thus, PhaR can downregulate production of EPSI under conditions that do not favor PHB synthesis. Moreover, a moderate repression by PhaR of the *wgaA* promoter, which drives the expression of ten *wga* genes involved in EPSII biosynthesis, was also demonstrated (Fig. 7C). As in *S. meliloti*, in *C. necator*, EPS and PHB production levels correlated either partially or completely, depending on the specific medium composition (J. Wang & Yu, 2007). In contrast, in the plant-associated nitrogen fixer *Azospirillum brasilense*, mutation of *phbC* led to approximately a 2-fold increase in EPS and KPS production. Notably, although a *phaR* homolog is annotated in the genome of *A. brasiliense*, it has not been functionally studied. Similarly, the *phbC* mutant of the extremophilic bacterium *Pseudomonas extremaustralis* produced more EPS than the wild type (Tribelli & López, 2011). Thus, the coregulation of EPS production and PHB biosynthesis might be not strictly conserved among PHB producing species. In *S. meliloti*, PHB synthesis could serve as an indicator of carbon homeostasis, signaling carbon excess relative to other nutrients. In species with unlinked regulation, EPS and PHB biosynthesis may compete for carbon substrates.

### PhaR targets multiple control points in central metabolism in response to C availability

The proteomic analysis revealed that PhaR has a broader role in modulating metabolism beyond regulating PHB and EPS metabolic branches. A key control point influenced by PhaR appears to be the regulation of pyruvate fate, in agreement with prior findings in *R. etli* (Dunn et al., 2002). In *R. etli*, mutation of *phbC* leads to elevated NAD(P)H and increased organic acid excretion, alongside enhanced activities of Pdh (pyruvate dehydrogenase), GltA (citrate synthase), Pyc (pyruvate carboxylase) and PckA (e.g., phosphoenolpyruvate carboxykinase). Dunn and collaborators proposed that the accumulation of redox power resulting from the inability to synthesize PHB in the *phbC* mutant post-translationally represses the activity of these key central metabolic steps. Our results, however, support the hypothesis that PhaR is a primary regulator of these enzymes. While PhaR activity as a transcriptional regulator is influenced by the presence of PHB, it operates in a PHB-independent manner. In the absence of PHB during active growth, PhaR favors the oxidative decarboxylation of pyruvate through upregulation of the pyruvate dehydrogenase complex (Pdh), while simultaneously repressing alternative conversion pathways by downregulating pyruvate phosphate dikinase (PpdK) and pyruvate carboxylase (Pyc). The activation of Bkd by PhaR may indirectly contribute to controlling PHB accumulation under specific growth conditions, as the degradation of branched-chain amino acids serves as a direct source of acetyl-CoA and redox equivalents. Moreover, the repression of citrate synthase (GltA) by PhaR suggests that limiting its activity may promote a greater allocation of acetyl-CoA and NADH generated from pyruvate decarboxylation toward PHB synthesis. In addition, PhaR might indirectly modulate C flux through the TCA cycle by regulating the IIA enzyme of the PTS phosphotransferase system, ManX. In *R. leguminosarum*, ManX controls the activity of TCA cycle enzymes by a yet unclear mechanism, likely involving protein-protein interactions (Sánchez-Cañizares et al., 2020). Derepression of *manX* under PHB-accumulating conditions could promote acetyl-CoA oxidation. Alternatively, higher levels of SMc01727 (a putative isocitrate lyase family protein) under these conditions may facilitate C shunting from acetyl-CoA through the glyoxylate cycle, providing intermediates to support EPS biosynthesis via an active gluconeogenic pathway. The role of PhaR sequestration in activating Tam, whose homologs have been shown to help alleviate the toxicity of *trans*-aconitate formed when the *cis*-isomer TCA-cycle intermediate accumulates (Cai et al., 2001), should be further investigated.

Previous studies reported that *phbC* mutants of *S. meliloti* strain Rm41 and *R. etli* were unable to grow on pyruvate, and on pyruvate or glucose as the sole carbon sources, respectively. Nevertheless, the corresponding double *phbC phaR* mutants regained this ability (Encarnación et al., 2002a; Povolo & Casella, 2009b). The repression of the gluconeogenic gene *ppdK* and the initial steps of the Entner-Doudoroff pathway by PhaR (Table 2, Fig. 7) provides a plausible explanation for the growth deficiencies observed in the *phbC* mutants. Activation of the upper segment of the ED pathway might not necessarily indicate that PhaR promotes the net catabolism of hexoses. Instead, it could facilitate a high C flux through a cyclic ED pathway, a mode of function previously reported in *S. meliloti* (Geddes & Oresnik, 2014).

We found no overlap between the PhaR regulon and the genes reported by D’Alessio and collaborators to exhibit altered expression in mutants defective for PHB biosynthesis grown under PHB-permissive conditions (D’Alessio et al., 2017). In *S. meliloti*, promoters containing a conserved binding motif for a Fnr-type transcriptional regulator (i.e., FixK proteins)— primarily located on the pSymA plasmid— are influenced by PHB accumulation under N-limiting, aerobic conditions (D’Alessio et al., 2017). In *B. diazoefficiens*, FixK_2_ operates within a regulatory cascade involving PhaR under microaerobic conditions (Quelas et al., 2016, 2024). Our findings suggest that, unlike in related rhizobia, the link between the expression of *fixK-*associated genes and the presence of PHB is not mediated by PhaR.

### PhaR’s influence may reach further than previously recognized

PhaR’s role may extend beyond its currently understood functions. Deeper phenotypic characterization is required to elucidate the significance of multiple regulated genes (Table 2), such as the genes encoding chemotaxis-related proteins McpT and McpZ (Baaziz et al., 2021), the manganese transport system substrate-binding protein SitA (Davies & Walker, 2007), the (S)-2-haloacid dehalogenase Dhe (Adamu et al., 2020), among others. Moreover, several strongly regulated targets, some of which have been confirmed to be directly controlled by PhaR at gene expression level, remain functionally uncharacterized. These include the small hypothetical proteins SMc03998 and SMc02051, the putative two-component sensor histidine-kinase SMb20609, the putative pilin glycosylation protein SMb21248, the conserved hypothetical proteins SMc01834 and SMb20727, making it difficult to fully understand the extent of PhaR-mediated regulation. In some cases, complex mechanisms might operate, where the regulation observed at the transcriptional level inversely correlates with the net effect measured at the protein level (e.g., the sugar ABC transporter ATP-binding protein SMc03815).

### PhaR and its role during symbiosis

Mutation of *phaR* in Sm2011 did not lead to any significant changes in symbiotic performance, in contrast to the results observed for *S. meliloti* Rm41 (Povolo & Casella, 2000a), where *phaR* mutation resulted in compromised symbiosis with the host plant *Medicago sativa*. The nodule number and plant dry weight increased in plants inoculated with Rm41 *phaR* mutant, although big part of the nodules appeared colorless and acetylene reduction activity was dramatically diminished (Povolo & Casella, 2000a).

Given the critical role of EPS production during rhizobial infection of the host plant, PHB accumulation in the N-limiting, C-rich rhizosphere may not merely fuel bacterial division and progression along the infection thread, as has long been postulated. Instead, it may indirectly activate EPS biosynthesis through PhaR sequestration, thereby influencing rhizobial competitiveness rather than infectivity. Regulation by *phaR* was also observed in nodule bacteria and bacteroids, as evidenced by the analysis of PhaP1-EGFP fluorescence in fresh nodule sections. This indicates that *phaR* is expressed and functions as a transcriptional repressor, even during the symbiotic differentiation of the bacteria. Under laboratory conditions, overexpression of *phaP1* and deregulation of other PhaR target targets in vegetative and symbiotic bacteria did not significantly affect infection or nitrogen fixation. However, in the natural environment, where constant inter- and intra-species competition occurs, unadjusted metabolism in the absence of PhaR would likely impair bacterial fitness, negatively affecting overall symbiotic efficiency.

### An integrated model of PhaR’s dual role as metabolic sensor and regulator

Figure 9 provides schematic representation of PhaR’s regulon mapped onto the metabolic pathways. PHB accumulation triggered by non-C nutrient limitation promotes a global proteome adjustment through sequestration of the transcriptional regulator PhaR to the granule surface. This sequestration induces the expression of the PhaR-direct targets phasins and PHB depolymerases, enabling controlled PHB accumulation. Although PHB biosynthetic genes are not directly repressed, the production of PHB precursors, such as acetyl-CoA and NAD(P)H, is likely reduced through the deactivation of their primary metabolic sources, including pyruvate dehydroganase and branched-chain amino acid degradation. In this context, an increased gluconeogenic flux seems required to offer pyruvate alternative fates other than its oxidative decarboxylation. Notably, gluconeogenesis may channel C toward the synthesis of alternative storage polymers, such as EPS, whose biosynthetic pathway is directly activated by PhaR sequestration. Furthermore, acetyl-CoA generated by PhaR-promoted turnover of PHB might be spared from complete oxidation. It is tempting to speculate that the direct activation of citrate synthase and the indirect regulation of TCA cycle-enzymes via ManX contribute to balancing fluxes between the TCA cycle and glyoxylate shunt. Metabolic flux analyses will be key to addressing this.

**Figure 9.**
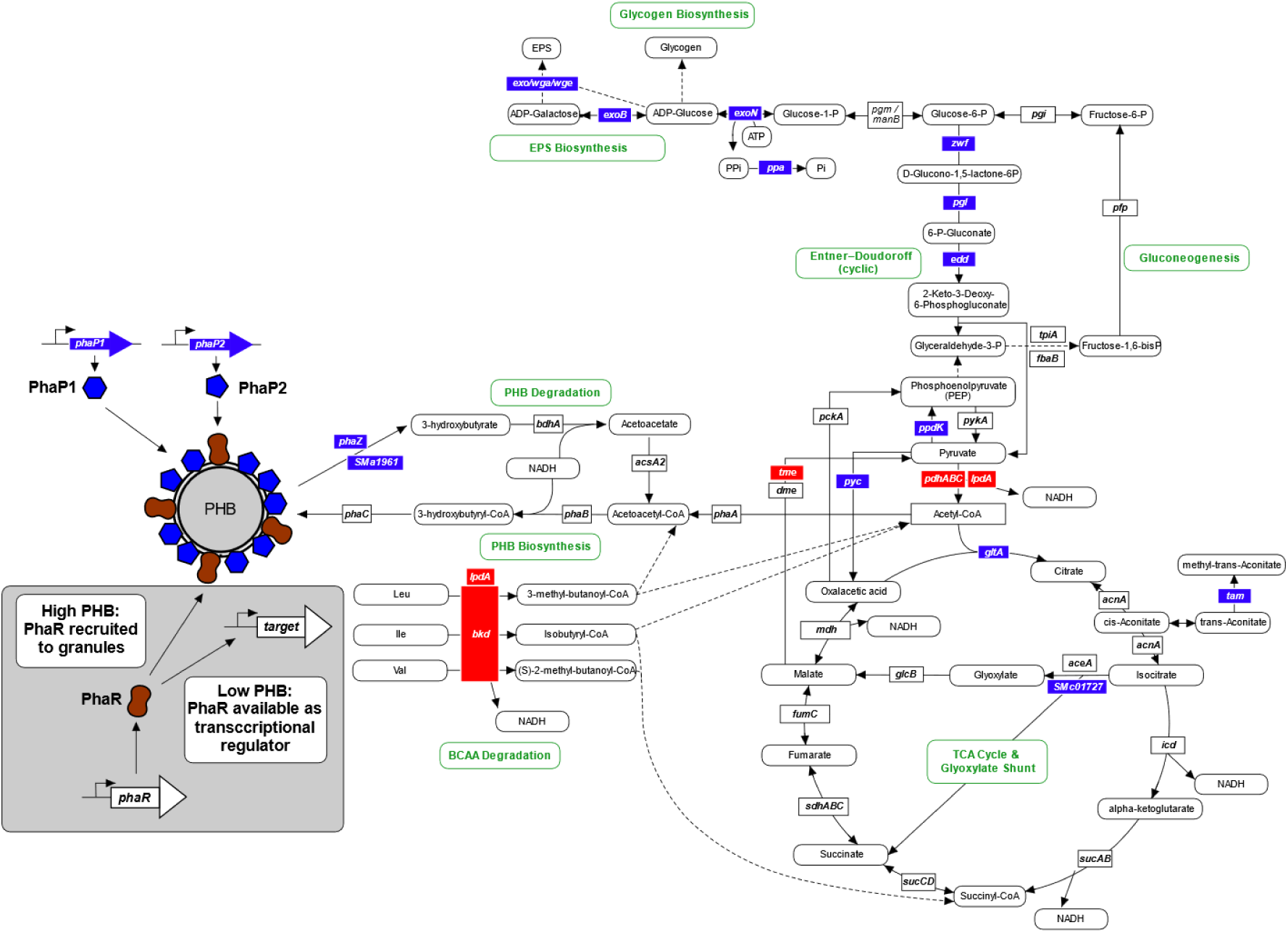
Proposed model of PhaR and PHB-dependent gene regulation in *S. meliloti*. Genes repressed by PhaR are shown in blue, and those activated by PhaR are shown in red. The names of major metabolic pathways are depicted in green. Rounded rectangles represent metabolites, while squared rectangles indicate enzyme names. Dashed lines represent multiple metabolic reactions, while solid lines represent a single reaction.

