## Supplementary figures and images for "A systems-level insight into PHB-driven metabolic adaptation orchestrated by the PHB-binding transcriptional regulator AniA (PhaR)"

### Figure S1

#### Figure S1

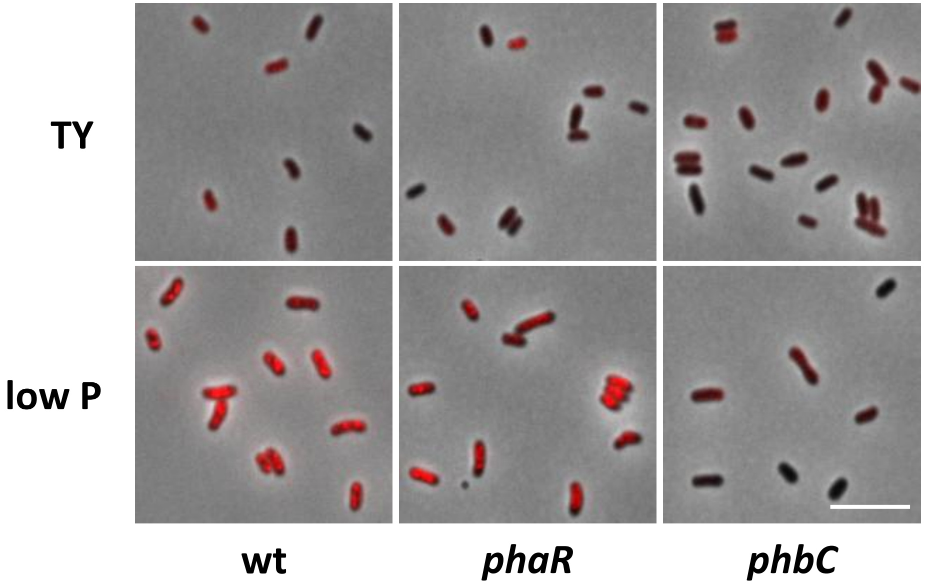
